# Functional profiling of extracellular vesicles from boar reproductive fluids

**DOI:** 10.1101/2025.10.11.681790

**Authors:** Veronika Kraus, Barbora Doleckova, Michaela Frolikova, Ondrej Sanovec, Daniela Spevakova, Zuzana Pilsova, Aneta Pilsova, Katerina Komrskova, Ondrej Simonik, Pavla Postlerova

## Abstract

Extracellular vesicles (EVs) are small membrane-bound structures that facilitate intercellular communication in the reproductive system, modulating gamete maturation, capacitation, immunomodulation, and fertilization. Despite pigs’ high relevance as a biomedical model, many aspects of EV biology remain poorly understood. EVs from boar seminal plasma (SP) are relatively well studied, whereas epididymosomes remain largely uncharacterized. Therefore, this study aimed to isolate and characterize EVs from the caput, corpus, and cauda regions of boar epididymis, as well as from SP, with their further precise analysis towards their interaction with sperm. We successfully obtained EVs from all studied fluids with sufficient purity. Importantly, our isolation protocol preserved the EVs’ ability to interact with sperm, demonstrated by staining with lipophilic dyes and biotin labeling experiments, confirming precisely their interaction and cargo transfer to sperm cells. Well-established EV markers, such as Alix and tetraspanins, were detected in the EVs, and additionally, phosphorylated, ubiquitinated, and sialylated proteins were uniquely identified. Furthermore, we employed a proteomic approach to characterize EV proteins and investigate their functional roles using the Gene Ontology (GO) database. This study contributes valuable insights into the molecular composition and functional properties of EVs from the male reproductive tract. It may provide a solid framework for further basic and translational research in reproductive biology and biomedicine.

## Introduction

Pigs are among the most important livestock species and, due to their high similarity to humans, represent a valuable biomedical model that is also highly relevant for the study of reproduction. Therefore, assisted reproductive technologies (ART) are a valuable approach to maintaining livestock breeding. However, in the case of the pig, some limitations make ART for this model less effective compared to other species (Dyck et al., 2014). Hence, improving ART outcomes in pigs is crucial. One potential approach to enhance these technologies involves modifying the media by adding components naturally present in reproductive fluids, such as small membrane structures known as extracellular vesicles (EVs) (Fazeli and Godakumara, 2024; Rodriguez-Martinez et al., 2024). EVs released by reproductive tract cells mediate molecular communication and deliver cargo between these cells and gametes (Tamessar et al., 2021; Gurunathan et al., 2022). Their cargo includes a variety of bioactive molecules and macromolecules, such as lipids, nucleic acids, and proteins, which influence key biochemical processes involved in gamete maturation, response of the immune system, or fertilization (Abels and Breakefield, 2016; Tamessar et al., 2021). The involvement of EVs in sperm life begins in the testes. Studies in mammals have shown that testicular EVs, also referred to as testisomes (Antalíková et al., 2022), are not only involved in intercellular communication but also likely participate in key processes such as spermatogenesis and steroidogenesis (Yun et al., 2019, Choy et al., 2022, Ma et al., 2023) A study by Antalikova et al. (2022) suggested that testisomes contain tetraspanin-enriched microdomains, including molecules such as CD9, CD81, and integrin alpha V. Following this initial interaction within the testes, the sperm come into additional contact with the EVs in the epididymis, known as epididymosomes. These EVs are secreted by epididymal epithelial cells (Rimmer et al., 2021; Tamessar et al., 2021; Barrachina et al., 2022). The epididymis, divided into caput, corpus, and cauda, provides a specialized environment for sperm maturation and storage. The apical blebbing, reflecting the ability of epithelial cells to release EVs, has been proposed as a marker of the mature phenotype of the epididymis (Hughes and Berger, 2015). As sperm transit through these regions, they are immersed in epididymal fluid, which contains secretions from the surrounding epithelial cells. Since the different parts of the epididymis are unique in their function and structure, it is to be expected that the epididymosomes represent a heterogeneous population of EVs with a cargo composition that is specific to their region of origin (Frenette et al., 2006; Domeniconi et al., 2016). This cargo diversity reflects the dynamic and region-specific roles epididymosomes play in sperm maturation and functional modulation, including protection of sperm during their storage in the cauda epididymis (Sullivan, 2015; Nixon et al., 2019; Tamessar et al., 2021). Epididymosomes also contribute to remodelling of the sperm plasma membrane through protein transfer, with tetraspanin CD9 identified in the bull model as a key mediator of sperm–EVs interactions (Nixon et al., 2019; Caballero et al., 2013). After the release of sperm from the epididymis, they encounter seminal plasma (SP) during ejaculation. This represents the final major exposure of spermatozoa to a heterogeneous population of EVs derived from the male reproductive organs. SP is the most significant source of EVs that originate from various tissues involved in SP production, most notably the bulbourethral glands, seminal vesicles, and prostate (Jonáková et al., 2007; Caballero et al., 2008; Kupcova Skalnikova et al., 2019). SP also contains epididymal secretions, and in pigs, epididymosomes have mainly been inferred from their putative presence in SP, with CD44 protein suggested as a potential marker (Alvarez-Rodriguez et al., 2019; Rodriguez-Martinez et al., 2024). However, specific details regarding epididymosomes in pigs remain unknown. EVs from SP, called seminal EVs, play a significant role in fertility by modulating sperm motility and capacitation, as well as by influencing the female reproductive tract during sperm transit (Piehl et al., 2013; Guo et al., 2019; Murdica et al., 2019; Vickram et al., 2021; Roca et al., 2022). Beyond their physiological functions, EVs are also implicated in pathological processes, including cancer progression and infections (Théry et al., 2009; Kalluri and LeBleu, 2020). Therefore, understanding the molecular composition of EVs, their mechanisms of binding to target cells, and the processes governing cargo sorting is essential.

The present study uniquely targets boar epididymosomes from the individual part of the epididymis (caput, corpus, and cauda), and EVs from SP. Special attention is given to the proteomic profiling of these EVs and their interactions with spermatozoa. EVs have been detected across different species, and research on porcine EVs has predominantly focused on those derived from SP. In contrast, studies on boar epididymosomes remain limited. Therefore, investigating epididymosomes in pigs holds the potential to provide novel insights into sperm maturation and fertilization processes. Such knowledge could contribute to improving biotechnologies related to *in vitro* fertilization and enhancing livestock reproductive efficiency.

## Materials and Methods

All chemical reagents were obtained from Sigma-Aldrich (St. Louis, USA) unless otherwise stated.

### Boar reproductive fluid and sperm collection

Semen collected from fertile Duroc boars bred for commercial artificial insemination was provided by the Insemination Station Skrsin (LIPRA PORK a.s., Rovensko pod Troskami, Czech Republic). Handling and care of animals were in accordance with Council Directive 98/58/EC, Act No. 154/2000 Coll., and Act No. 246/1992 Coll. of the Czech National Council. The male reproductive tissues were obtained from sexually mature Prestice Black-Pied boars, 8 to 12 months old, from the Biofarm Sasov (Jihlava, Czech Republic). Organs were transported on ice and processed immediately. Tissues were collected post-mortem under veterinary supervision in compliance with Directive 2010/63/EU and Czech regulations (208/2004 Sb.). No procedures were performed on live animals.

Small tissue pieces from the caput, corpus, and cauda regions of the epididymal ducts were incubated in phosphate-buffered saline (PBS; P4417) at 38 °C and 5 % CO₂ for 30 min. The resulting suspension was subsequently filtered and centrifuged at 500 × g for 10 min to exclude spermatozoa. The ejaculate was centrifuged at 300 × g for 10 min for sperm separation. Spermatozoa from the caput and cauda epididymis were subsequently washed three times in PBS by centrifugation at 500 × g for 10 min and used for functional studies with EVs. Alternatively, pellets of 5 × 10^7^ /ml were used for protein isolation. The fluids obtained from the caput, corpus, and cauda epididymis and SP were used for the isolation of EVs.

### Isolation of EVs

EVs were isolated from frozen (−80 °C) epididymal fluid of caput, corpus, and cauda and SP in three main steps, as shown in Figure 1. Firstly, epididymal fluid and seminal plasma were thawed in a water bath at 37 °C. After thawing, the fluids were first centrifuged at 300 × g for 10 min. The supernatant was transferred to a clean tube and centrifuged at 2,000 × g for 10 min at 4 °C. The supernatant was transferred again to a clean tube and centrifuged at 10,000 × g for 20 min at 4 °C. If the obtained supernatant exceeded a volume of 2 ml, it was subsequently concentrated using a Pierce^TM^ protein concentrator PES, 100K MWCO, 5-20 ml (Thermo Fisher Scientific, Waltham, USA) to a maximal volume of 2 ml. The concentrated EVs were purified from proteins using pure-EVs: Size Exclusion Chromatography (SEC) columns (HansaBioMed, Tallinn, Estonia). Before loading the concentrated samples, the columns were first washed with 30 ml of sterile and filtered PBS buffer. After washing, a concentrated sample of epididymal fluid or seminal plasma was loaded into the column and collected in 19 fractions of 500 μl. Once the sample was loaded into the gel matrix, additional sterile and filtered PBS was applied to the column to prevent drying. After the 19 fractions were collected, the column was washed with 20 ml of sterile and filtered PBS buffer and stored at 4 °C for further use. Fractions 6–12, containing lower protein concentrations, were selected for further analyses based on protein concentration measurements (see Chapter Protein Concentration Measurement). From fraction 13 onwards, protein concentrations increased, and therefore, the fractions were not used for further analyses of EVs. Fractions that were not immediately used were stored at −80 °C with 0.25 M trehalose.

**Figure 1.**
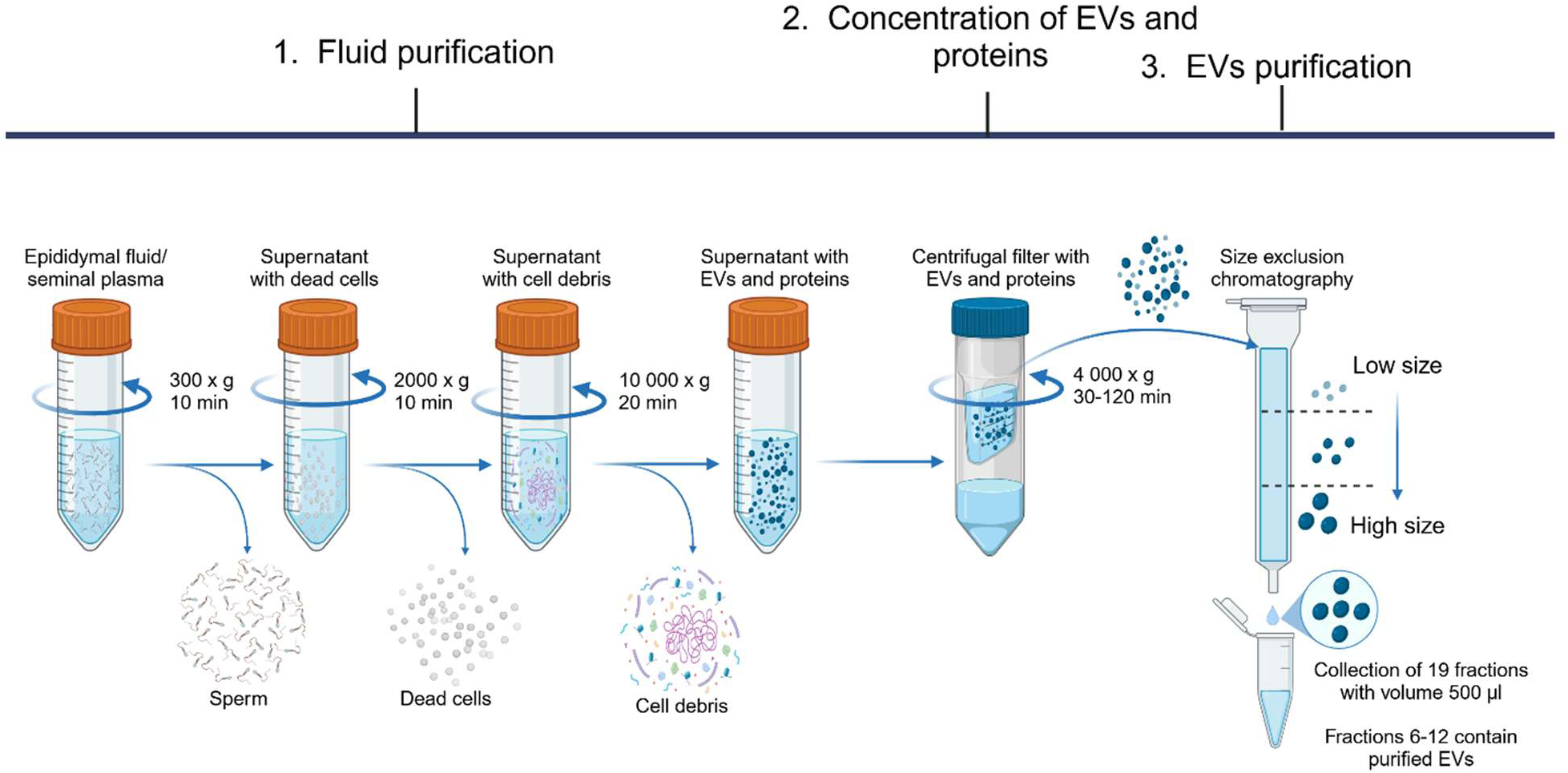
Isolation of EVs. Firstly, biofluids were purified from cells, debris, and macrovesicles through differential centrifugation. Consequently, the fluids were concentrated into a smaller volume, which was then transferred to size exclusion columns (SEC), where particles were separated according to their size. The outcome consisted of 19 fractions, each with a volume of 500 µl. Based on further characterization, fractions 1–5 contained only PBS, fractions 6–12 comprised purified EVs, and fractions 13–19 contained EVs along with progressively increasing concentrations of co-isolated proteins. (Created with BioRender.com)

The experimental design comprised three main aims: isolation and identification of EVs, proteomic analysis, and functional studies. A combination of SEC, dynamic light scattering (DLS), and electron microscopy (EM) was employed for the identification and characterization of EVs. Proteomic analysis involved SDS-PAGE, western blotting, and mass spectrometry (MS), while functional studies employed CASA system (Computer-Assisted Sperm Analysis), lipophilic staining, biotin labeling, and microscopic techniques (confocal, super-resolution, and electron microscopy) (Figure 2).

**Figure 2.**
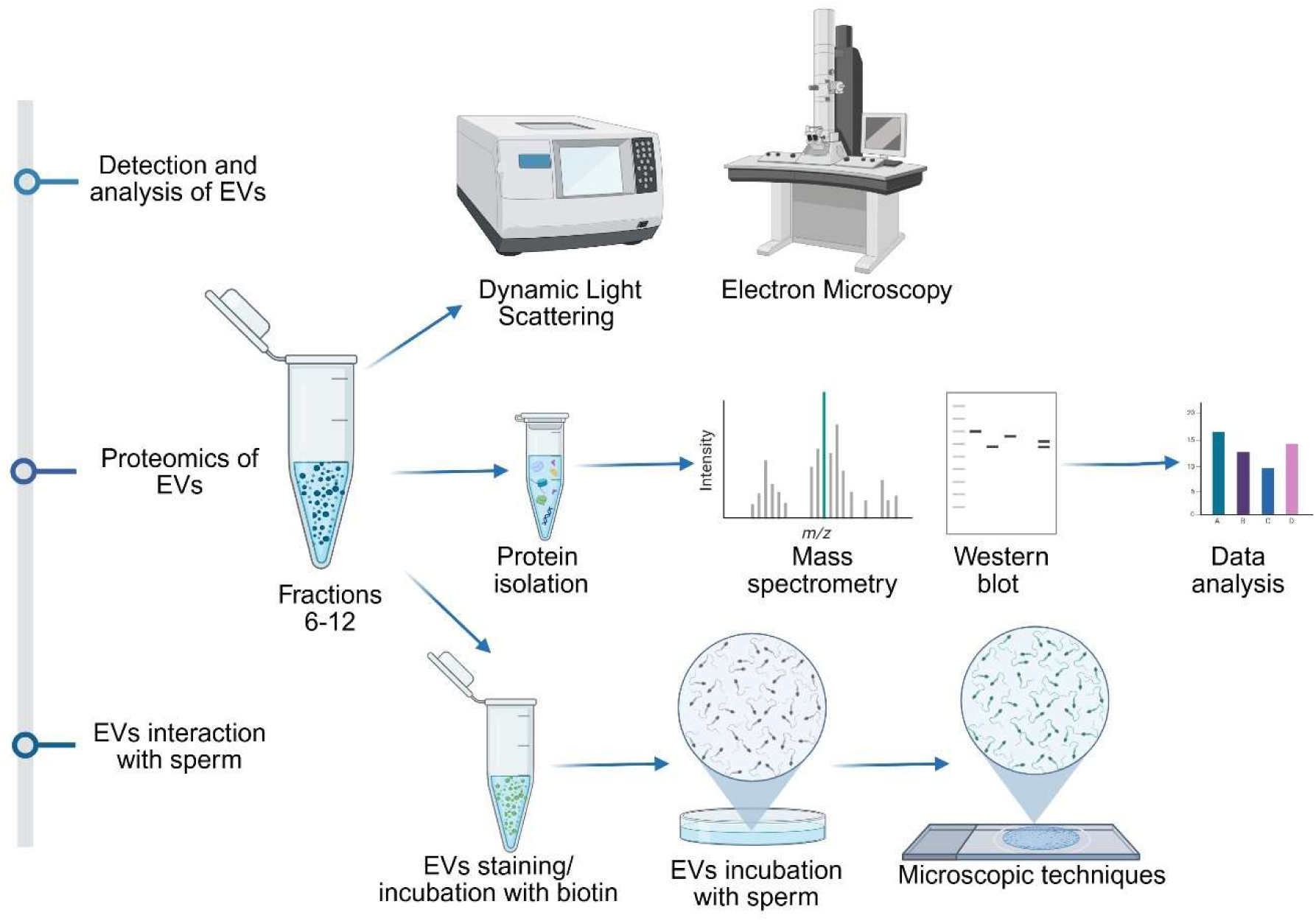
Experimental design for studying EVs. EVs isolated from epididymal fluid and SP were characterized by DLS and electron microscopy, followed by proteomic analysis using Western blot and mass spectrometry. Functional assays were then performed to evaluate EVs’ interactions via dual staining with lipophilic dyes and cargo transfer through biotinylation labeling. (Created with BioRender.com)

### Protein concentration measurement

Protein concentration was measured in all fractions using the Pierce^TM^ BCA Protein Assay Kit (Thermo Fisher Scientific). First, the prepared standards were diluted in PBS according to the datasheet to the appropriate concentrations Subsequently, a working solution diluted 50:1 was prepared from BCA reagent A and BCA reagent B. Triplicates of 25 μl from both standards and samples were loaded on a 96-well plate, and 200 μl of the prepared working solution was added to this followed by mixing and incubation at 37 °C for 30 min. After incubation, the intensity of the colorimetric staining was measured at 562 nm by a microplate reader (Tecan, Grödig, Austria).

### Dynamic Light Scattering

DLS measured the distribution of EVs in individual fractions (6-12). The EVs were measured at 25°C, at a wavelength of 633 nm, and a scattering collection angle of 90°. They were measured in a microcuvette ZEN2112 with a volume of 50 μl using a Zetasizer Nano ZS90 (Malvern Panalytical, Malvern, UK).

### Electron Microscopy (EM)

To verify the presence of EVs in fractions isolated from epididymis and seminal plasma, samples were examined using Transmission Electron Microscopy (TEM). For negative stain analysis, 4 μl of each sample was applied to a glow-discharged TEM copper grid with a continuous layer of carbon (300 mesh) and incubated for 1 min. The grid was then blotted using filter paper, successively washed in three drops of demineralized water, and stained with 2% uranyl acetate (Electron Microscopy Sciences, Hatfield, USA) (aq., 2 drops each for 30 s) and allowed to air dry.

Cryo-electron microscopy (Cryo-TEM) was employed to visualize the interaction between EVs and sperm after a biotinylated experiment. A 4 μl of sperm + EVs sample was applied to a glow-discharged copper cryo-EM grid (Quantifoil R 2/1, 300 mesh) and incubated for 1 min in a Leica GP2 Plunge Freezer. The grid was then automatically blotted using filter paper for 4 s and immediately vitrified by plunging into liquid ethane cooled to –180 °C. After vitrification, the sample was transferred to a dual-grid cryo-holder from Simple Origin and inserted into the transmission electron microscope (TEM).

Electron microscopy data were acquired in a semi-automated mode using SerialEM software (Mastronarde, 2005) on a Jeol JEM-2100Plus TEM microscope equipped with a LaB6 electron gun and a TVIPS XF416 CMOS camera, operated at 200 kV. For cryo-TEM analysis, overview images were acquired at x8k (pixel size of 1.44 nm) and details were collected at x20k magnification (pixel size of 0.59 nm) with a 3-second exposure time and a total electron dose of 50 electrons/Å².

### Labeling Strategies for Extracellular Vesicles

#### Deproteination of EVs by proteinase K (Prot K)

From the fractions with pure EVs (6–12 for epididymal fluid and 6–11 for SP), 100 μl of supernatant was collected and mixed. From the supernatant, 500 μl was transferred to a new tube, and Prot K (20 mg/ml) for Total Exosome Isolation (from plasma) (Invitrogen, Thermo Fisher Scientific) was added at a final concentration of 50 μg/ml. Samples with and without Prot K were incubated at 37 °C for 30 min with stirring. Subsequently, c*O*mplete Mini protease inhibitors (Roche, Basel, Switzerland) were added to the samples, and the samples were incubated on ice for 30 min with periodic vortexing every 10 min. Subsequently, deproteinated samples were purified by SEC in the same manner as was described above.

### Double lipophilic staining protocol for sperm-EVs detection

EVs from the cauda epididymis and SP were stained using CellBrite 480 lipophilic dye (30090-T, Biotium, Fremont, USA) diluted 1000× directly into a prepared mixture of isolated EVs from fractions 6–12 (total volume 100 µl). The EVs’ mixture was incubated in the dark for 15 min at 37 °C. After incubation, PBS was added to each group at a 1:2 ratio to wash off the dye. Samples were transferred into 2ml Amicon Ultra filters (100 kDa) (UFC210024, Merck Millipore, Burlington, USA) and centrifuged at 4,500 × g until a reduction of volume to 200 µl, and then supplemented with PBS to 600 µl.

Sperm were washed twice in prepared and heated sperm-TALP medium (Tyrode’s Albumin Lactate Pyruvate; 114 mM NaCl, 3.2 mM KCl, 0.3 mM NaH_2_PO_4_.H_2_O, 10 mM sodium lactate, 2 mM CaCl_2_.2H_2_O, 0.5 mM MgCl_2_.H_2_O, 10 mM HEPES, 25 mM NaHCO_3_, 0.6 % BSA, 1 mM sodium pyruvate) and centrifuged at 300 × g for 7 min at room temperature (RT). The initial sperm concentration of 2.5 × 10^8^ was set equally for all experimental groups. Washed spermatozoa were further incubated with CellBrite 640 lipophilic dye (#30090-T, Biotium, Fremont, USA) in the same manner as EVs and washed again twice in 1 ml of sperm-TALP medium. After EVs and sperm staining, green-stained EVs were co-incubated with red-stained spermatozoa in the dark at 37 °C for 90 min (Figure 3a).

**Figure 3.**
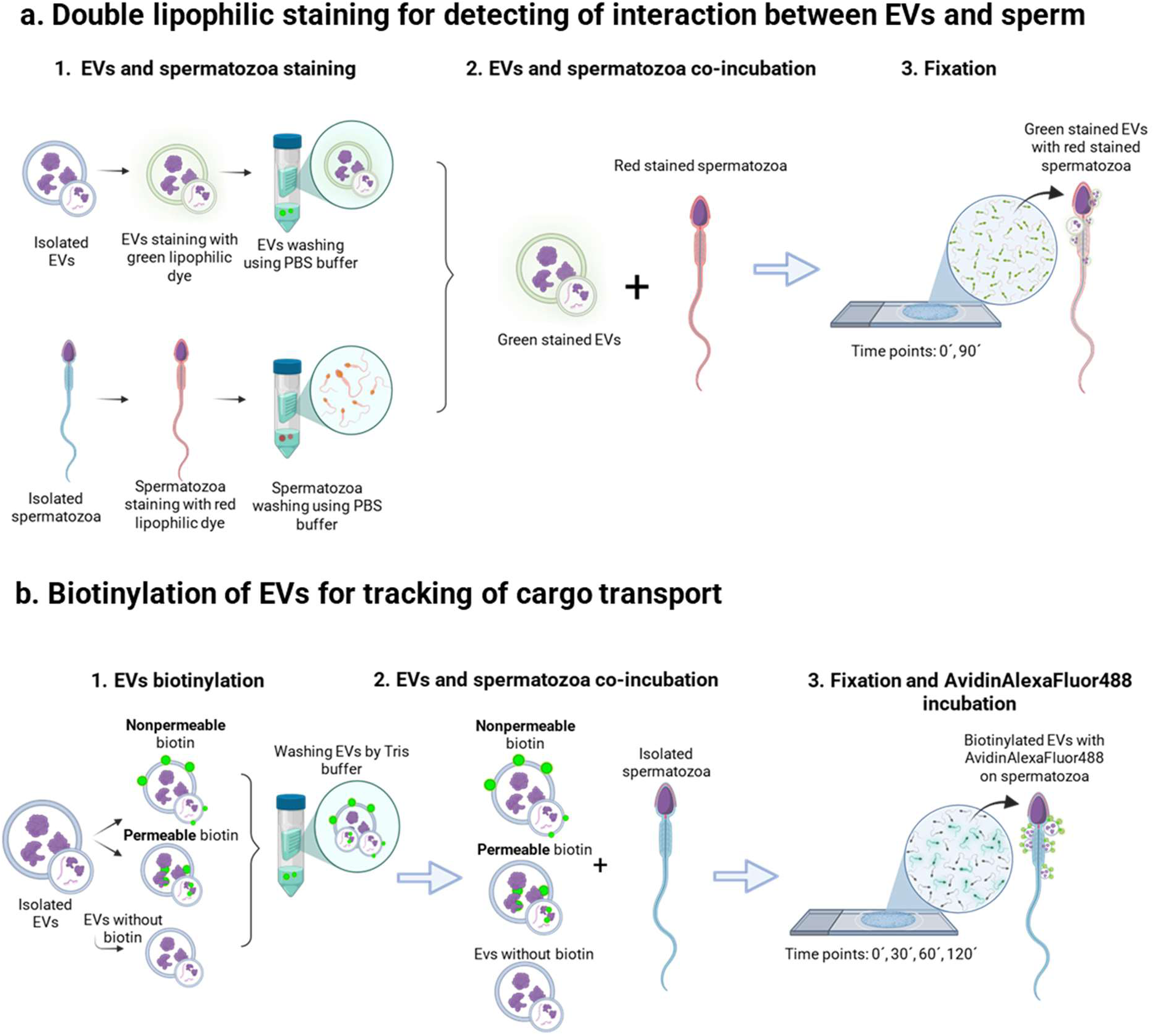
Labeling and visualization of EV-sperm interactions. To monitor the interaction between EVs and spermatozoa, two complementary labeling approaches were used. First, double lipophilic staining was applied to visualize the physical interaction between green-labeled EVs and red-labeled spermatozoa. Co-localization of both fluorescent signals indicated direct contact between vesicles and sperm. In parallel, biotin labeling was used to assess the cargo transfer from EVs to sperm. EVs were labeled either with permeable or non-permeable biotin, allowing for the detection of the EVs’ cargo internalized within sperm cells. (Created with BioRender.com)

### Biotinylation of EVs for the detection of cargo transport

The EVs’ biotinylation was performed and modified according to Zhou et al. (2019). Permeable biotin (EZ-Link™ BMCC-Biotin, 21900, Thermo Fisher Scientific) was dissolved at 0.5 mg/100 µl in 99.8 % dimethylformamide (100379, Millipore) and non-permeable biotin (Sulfo-NHS-SS-Biotin, 21331, Thermo Fisher Scientific) at 1 mg/100 µl in PBS. EVs isolated from gradient fractions 6–12 were pooled and adjusted to a final volume of 300 µl per sample. Three experimental groups were prepared and incubated for 1.5 hours at RT: the first group was incubated with permeable biotin at a final concentration of 1.34 mM, the second group with non-permeable biotin at a final concentration of 0.63 mM, and the third was a control group with pure EVs without biotin. PBS purified on 20 ml, 100 kDa filters (88533, Thermo Fisher Scientific) was added to each group to achieve the same input volume (1 ml). At the end of the incubation period, 25 mM Tris-HCl buffer (pH 8) was added to each group in a 1:2 ratio. The samples were centrifuged on 2 ml Amicon Ultra filters (100 kDa) at 4,500 × g, until the volume was reduced to 200 µl, and then 600 μl of filtrated PBS was added to the prepared samples. Spermatozoa were washed twice in the prepared sperm-TALP medium and centrifuged at 300 × g for 7 min at RT. The initial spermatozoa concentration of 2.5 × 10^8^ was set equally for all experimental groups. Biotinylated EVs were co-incubated with spermatozoa in the dark at 37 °C for 90 min (Figure 3b). Supporting data on sperm motility were acquired during this experimental part, and total and progressive motility were measured by Computer Assisted Sperm Analysis (CASA) (ISAS, Proiser, Valencia, Spain).

### Fixation on slides

A 50 μl of sperm samples incubated with EVs from double lipophilic staining were collected after 90 min, and biotin labeling experiments were collected every 30 min and immediately fixed with 50 µl of 3.2 % para-formaldehyde (PFA; 047377, Thermo Fisher Scientific) onto polylysine-coated coverslips (8000104, Hirschmann, Germany). After incubation for 15 min at RT, samples were rinsed in PBS twice. The samples from the biotin-labeling experiment were incubated with Avidin AlexaFluor488 (SA-5001, Thermo Fisher Scientific) diluted 1:300 in PermWash (BDB557885, Thermo Fisher Scientific) and washed twice in PBS. The rinsed slides from both experiments were stained with DAPI solution (0.85 μg/ml, Thermo Fisher Scientific) and incubated at RT, in the dark for 10 min. Another wash was then performed, followed by a final immersion of the slide in deionized water (dH_2_O), and the slide was left to dry. After drying, the samples on coverslips were mounted on slides (631-1553, Avantor, Radnor, USA) with 10 µl of ADVI S mounting medium (ADM-005, Life M, Ricany u Prahy, Czech Republic). The samples were imaged as Z-stacks on a confocal microscope LSM 880 (Carl Zeiss, Oberkochen, Germany). Images from dual-stained samples were deconvolved using Huygens Professional (version 25.04) software, followed by surface reconstruction (rendering) in Imaris software (version 9.9.1). The sperm plasma membrane was processed using Surface function with surface material set to be transparent. EVs were processed in two ways. First, the Spot function was used to visualize the localization and size/signal intensity of EVs. Secondly, the rendering of EVs was performed using the Surface function. Objects corresponding to EVs were color-coded based on the shortest distance from the sperm plasma membrane. The DAPI signal was left in its native form.

### Proteomics

#### Isolation of proteins from EVs

A supernatant (100 μl) was taken from fractions containing pure EVs, mixed, and the resulting volume was concentrated to 100 μl using Amicon Ultra 100 K centrifugation filters. Subsequently, 1:1 Pierce^TM^ RIPA buffer (Thermo Fisher Scientific) was added to the centrifuged EVs, and the entire supernatant with EVs was transferred into a clean tube. To the total volume of EVs with RIPA, 1:5 Bolt^TM^ LDS Sample Buffer (4x) was added, and samples were sonicated on ice by sonicator (Qsonica, LLC, Newtown, CT, USA; amplitude 25 %, impulse 6×10 s, break on ice 20 s). Finally, all protein samples were reduced with 5% mercaptoethanol and incubated at 95°C for 5 min, except those derived from biotinylated extracellular vesicles, which were processed under non-reducing conditions.

#### Isolation of proteins from sperm

A pellet of boar sperm or sperm with EVs were incubated for 30 min at 4 °C, and subsequently boiled for 5 min at 95 °C in 2× concentrated non-reducing Laemmli sample buffer (20 % (v/v) glycerol; 4 % (w/v) Sodium Dodecyl Sulfate (SDS); 0.005 % (w/v) bromophenol blue; 0.125 M Tris-HCl, pH 6.8) with protease inhibitors. The samples were used for SDS-PAGE.

For the isolation of cytoplasmic and the rest of the pellet proteins from boar sperm incubated with EVs, the Subcellular Protein Fractionation Kit for Cultured cells (78840, Thermo Fisher Scientific) was used. Cytoplasmic extraction buffer (CEB) was added first, incubated with spermatozoa for 10 min at 4 °C, followed by centrifugation at 500 × g for 5 min at 4°C. The supernatant with cytoplasmic proteins was collected, and the Pellet Extraction Buffer (PEB) was added to the rest of the pellet. The sample was incubated for 10 min at RT and centrifuged at 16,000 × g for 5 min. The supernatant was transferred to a new tube, and the pellet was discarded. The 2× concentrated non-reducing Laemmli sample buffer was added to all tubes containing supernatant from different subcellular fractions in a ratio of 1:1. The samples were boiled for 5 min at 95 °C, and the protein fractions were subjected to SDS-PAGE.

### SDS-electrophoresis and Western Blot Analysis

For mass spectrometry analysis, samples containing isolated proteins from EVs were loaded onto a 4–20 % gradient gel (Bio-Rad, Hercules, USA) and were washed in dH_2_O and stained in 0.05 % (w/v) Coomassie Brilliant Blue R-250 solution (in 35 ml acetic acid; 250 ml ethanol; 215 ml dH2O) at RT overnight. The next day, the gels were incubated in the destaining solution (350 ml ethanol, 100 ml acetic acid, 550 ml dH_2_O) until the background was decolorized. Each sample lane (EVs from caput, corpus, cauda and SP) was cut into four sections based on molecular weight ranges (200–100 kDa, 100–50 kDa, 50–25 kDa, and 25–10 kDa) (Supplementary Figure S1), placed into 1 % (v/v) acetic acid and then subjected to mass spectrometry (MS) analysis. Firstly, protein bands were excised from SDS-PAGE gels and subjected to destaining in a solution of 25 mM ammonium bicarbonate (AMBIC) and 50 % acetonitrile (ACN) until clear. Following destaining, the gel pieces were dehydrated with 100 % ACN. Proteins were reduced in 100 mM dithiothreitol (DTT) at 60°C for 30 min, then washed sequentially with ultrapure water and ACN. Finally, they were alkylated with 100 mM iodoacetamide (IAA) in the dark for 30 min at 20°C. Additional washes in water and ACN removed excess IAA. To remove N-linked glycans, gel pieces were incubated with PNGase F (New England Biolabs) for 2 hours at 37 °C. After deglycosylation, the gel pieces were washed three times with alternating washes of water and ACN, dried in a SpeedVac, and digested overnight with Trypsin Gold (Promega) in 50 mM AMBIC at 37 °C. The resulting peptides were extracted by incubating the gel pieces with 100% ACN. ACN and the digestion supernatant were combined, dried by SpeedVac, and reconstituted in 0.1% formic acid for subsequent LC-MS/MS analysis. Peptide samples were loaded onto Evotip Pure tips (Evosep) according to the manufacturer’s instructions. Liquid chromatography was performed using the Evosep One system with the standard 30 samples/day method. Peptide separation was achieved on a reverse-phase PepSep C18 analytical column (15 cm × 150 µm, 1.5 µm; Bruker Daltonics). Mass spectrometric analysis was carried out on a timsTOF SCP instrument (Bruker Daltonics) operating in data-dependent acquisition (DDA) mode. The instrument was configured with an ion mobility range of 0.7–1.6 V·s/cm² and a m/z range of 100–1700. The acquisition cycle consisted of a TIMS survey scan (166 ms), followed by five parallel accumulation–serial fragmentation MS/MS scans, resulting in a total cycle time of 1.04 s. Raw data were analyzed using PEAKS Studio v12.5 (Bioinformatics Solutions Inc.) against the Sus scrofa (Uniprot UP000008227) databases. Variable modifications included methionine oxidation, N-terminal acetylation, and deamidation of asparagine and glutamine residues. The MS data are available upon request from the authors.

For Western blot (WB) analysis, the protein extracts were separated by 10 or 12 % SDS-PAGE (10 or 12 % (w/v) Acrylamide/Bis acrylamide solution (Bio-Rad); 1.5 M Tris-HCl (Bio-Rad), pH 8.8; 0.1 % (w/v) SDS; TEMED; 0.1 % (w/v) ammonium persulfate) and 4 % stacking gel (4 % (w/v) Acrylamide/Bis-Acrylamide solution; 0.5 M Tris-HCl pH 6.8 (Bio-Rad); 0.1 % SDS; TEMED; 0.1 % (w/v) ammonium persulfate), and the proteins were transferred to the PVDF membrane (Millipore). The molecular weight of the proteins was assigned by Precision Plus Protein Dual Color Standards (Bio-Rad). The membranes were blocked in 5% Blotto, non-fat dry milk (Santa Cruz Biotechnology, Dallas, USA) diluted in PBS and incubated with primary antibodies: anti-CD9/MRP1 (bs2489R, Bioss, Beijing, China), anti-CD81 (NBP2-20564, Novus Biologicals, Centennial, USA), anti-ALIX antibody (ab88388, Abcam, Cambridge, UK), anti-calnexin polyclonal antibody (10427-2-AP, ThermoFisher Scientific), anti-phosphoserine antibody (AB1603, Merck Millipore), anti-Ubiquitin FK2 (BML-PW8810, Enzo Biochem, Inc., Farmingdale, USA) diluted 1:250 in PBS for anti CD9 and anti-CD81 antibodies or 1:500 for anti-ALIX, anti-calnexin, anti-Phosphoserine or anti-ubiquitin antibodies and incubated at 4 °C overnight. After incubation, the membranes were washed in PBS and incubated with a secondary antibody, goat anti-rabbit IgG or goat anti-mouse IgG conjugated to horseradish peroxidase (Bio-Rad), diluted 1:3,000 in PBS for 1 hour at RT, and consequently washed in PBS. Sialic acids (Sia) were detected on PVDF membrane and blocked in 1 % (w/v) gelatine from cold water fish skin for 1 hour at RT. Afterwards, the membrane was incubated at 4 ◦C overnight with biotin-labeled *Sambucus nigra agglutinin* (SNA) lectin (B-1305, Vector Laboratories) diluted in HEPES buffer (10 mM HEPES; 0.1 mM CaCl_2_; 0.15 M NaCl; pH 7.5) to the final concentration of 1 μg/ml. Then, the membrane was washed in PBS and incubated with Avidin conjugated with Horseradish Peroxidase (avidin-HRP; A-3151) diluted 1:1000 in PBS for 1 hour at RT and consequently washed in PBS.

In a separate experiment, proteins isolated from biotinylated EVs and from sperm incubated with biotinylated EVs were transferred onto a low-fluorescence PVDF membrane (Bio-Rad). These membranes were blocked overnight at 4 °C with 1 % (w/v) gelatine, washed in PBS, and then incubated with avidin-HRP conjugate diluted 1:1000 in PBS for 1 hour at RT. All membranes were developed with SuperSignal™ Chemiluminescent Substrate (Thermo Fisher Scientific), and images were captured using the Azure c600 imaging system (Azure Biosystems, Inc., Dublin, USA). As a negative control, a membrane without primary antibodies or lectin was used. The membranes were stained with 0.05 % Coomassie brilliant blue R-250 solution for 5 min at RT and washed in a destaining solution to determine the total protein profile.

### Dot Blot analysis

The 5 μl of native biotinylated EVs, biotinylated EVs treated with Prot K, and non-biotinylated EVs were loaded onto a low-fluorescence PVDF membrane (Bio-Rad), and the membrane was allowed to dry for 30 min at RT. The membrane was then washed with PBS and blocked in 1% (w/v) gelatine overnight at 4 °C. The next day, the membrane was washed in PBS and incubated with avidin-HRP conjugate diluted 1:1000 in PBS for 1 hour at RT. The membranes were developed with SuperSignal™ Chemiluminescent Substrate (Thermo Scientific), and images were captured using the Azure c600 imaging system (Azure Biosystems, Inc.).

### Data analysis

The densitometric analysis was performed using the ImageJ software (Java-based image processing program, LOCI, University of Wisconsin, USA). Statistical analyses and the generation of Figures 1–3 were conducted with BioRender.com. To compare data between groups from the densitometric analysis, non-parametric statistical tests such as the Kruskal–Wallis test were used.

Data obtained from MS were analysed in Python (version 3.13.5) within the Spyder integrated development environment. *Pandas* (version 2.3.0) was used to load, organize, and preprocess the dataset, including protein list manipulation. *NumPy* (version 2.3.1) supported numerical operations and array-based data handling. Functional annotation and enrichment analysis were performed using *GProfiler* (version 1.0.0). Based on the list of identified proteins, Gene Ontology (GO) terms were analyzed. Enrichment was tested using a hypergeometric test, with multiple testing correction via the g:SCS method (Set Counts and Sizes). The complete *Sus scrofa* genome was used as the statistical background. GO analysis was divided into biological processes (BPs), Kyoto Encyclopedia of Genes and Genomes (KEGG) pathways, and cellular localization to enable a comprehensive analysis of the proteins. For more specific functional insights, these three categories were considered together as GO terms and visualized using a bubble plot. *Matplotlib* (version 3.10.3) and *Seaborn* (version 0.13.2) were applied to generate basic graphical representations of results and enabled advanced visualizations highlighting significantly enriched biological categories in an interpretable format. Visualization of unique and shared proteins across sample categories was done using Venny 2.1 (Oliveros, 2007-2015). A heatmap of the identified peptides was created using the BioRender platform (BioRender.com) for illustrative and presentation purposes. The stylistic editing and translation of certain sections of this manuscript were assisted by OpenAI’s ChatGPT (version GPT-4, July 2025), while the authors remain responsible for the final content.

## Results

### Detection and identification of EVs in fluids from the boar reproductive tract

EVs from fluids of the caput, corpus, and cauda regions of the epididymis, as well as from boar seminal plasma, were successfully isolated by combining centrifugation with SEC. SEC yielded 19 fractions, in which we measured protein concentration using the BCA assay and observed two peaks of protein concentration. The first peak increased in fractions 6 to 12, which likely correspond to proteins associated with EVs present in the fluids. The second peak, starting from fraction 13 and rising toward the final fraction, likely represents free proteins from the respective fluids (Figure 4a,b). Therefore, for subsequent experiments, we used fractions 6–12, which contain EVs and a lower concentration of free proteins. In the individual fractions (6–12), moderate heterogeneity of EVs was observed, with the highest polydispersity index indicating the broadest size variability in fraction 12 from the corpus fluid (0.53 ± 0.08), and the lowest in fraction 11 (0.22 ± 0.04) in the cauda fluid. We also observed a decreasing trend in the Z-average, which reflects the average hydrodynamic size of vesicles, across increasing fraction numbers in samples from the epididymal fluids. In contrast, samples from SP showed smaller variation in Z-average, though with a similar decreasing tendency. Among all fluids, the largest (307.1 ± 186.6 nm) EVs were detected in the corpus, and the smallest (113.4 ± 5.2 nm) EVs were detected in the caput, suggesting the presence of both microvesicles and exosomes in this region as well as in other fluids (Figure 4c). By pooling fractions 6–12, we obtained samples enriched in EVs, which were subsequently identified using TEM (Figure 4d). EVs appear as sharply defined spherical structures with varying electron densities (i.e., lighter and darker contrast) and visible membranes. Depending on the EVs’ population, some vesicles may appear damaged or deformed, for example, with ruptured membranes. EVs are often observed in clusters, and differences in their size, concentration, and background composition can be seen between individual fractions. The images also reveal variability in EVs’ morphology, size (typically ranging from 30 to 150 nm), and abundance across different regions of the epididymis and SP (Figure 4d).

**Figure 4.**
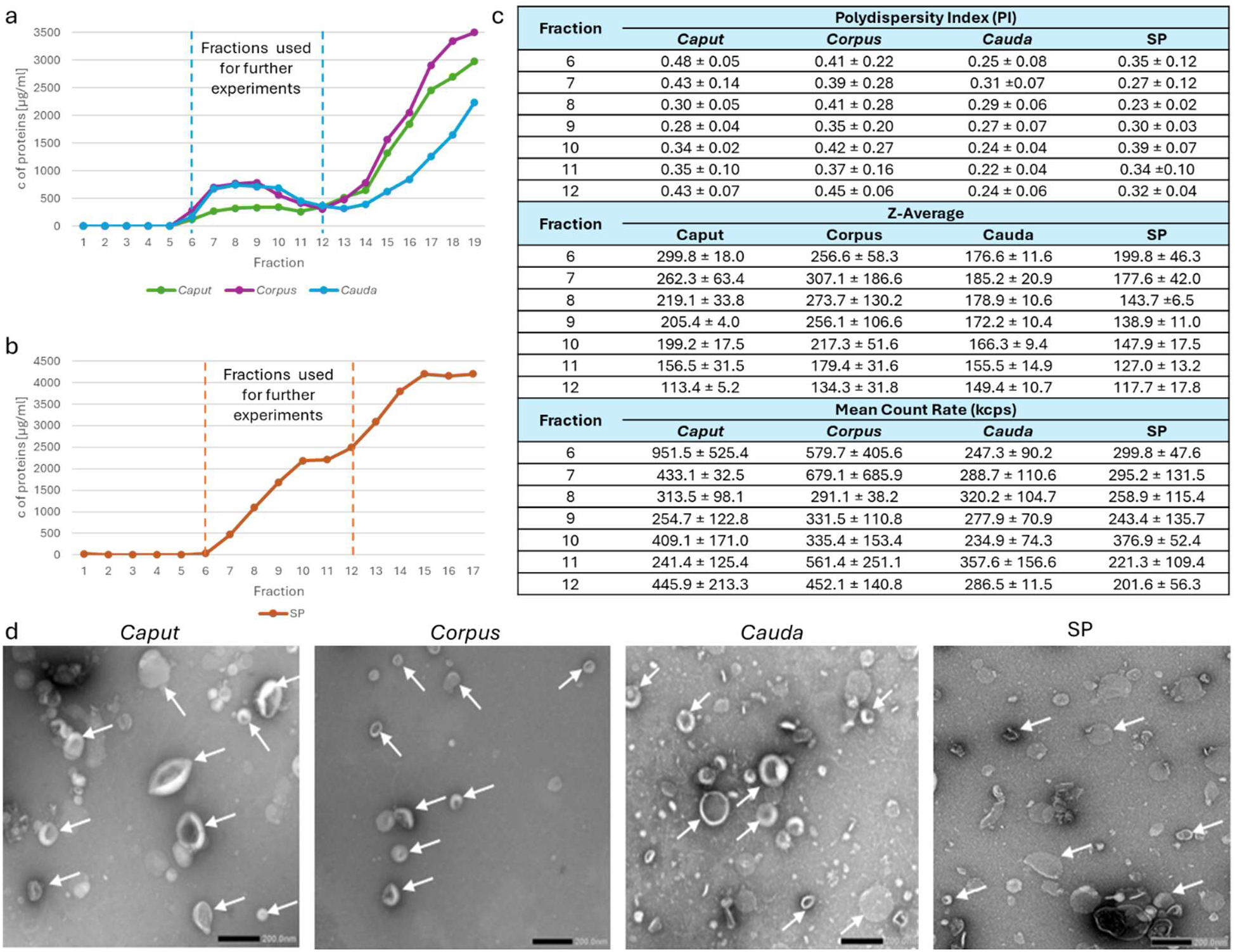
Detection and characterization of EVs in fluids from the male reproductive tract. **(a, b)** Protein concentrations measured in individual SEC fractions revealed two distinct peaks: the first in fractions 6– 12, corresponding to the most purified EV-containing fractions (indicated by a dashed line), and the second in fractions 13–19, likely representing free proteins. **(c)** Fractions 6–12 contained heterogeneous particles, as indicated by the polydispersity index (PI), with sizes corresponding to both exosomes and microvesicles (Z-average: 307.1 ± 186.6 to 113.4 ± 5.2 nm). The average number of photons detected per second during DLS analysis, referred to as the Mean Count Rate, is expressed in kilocounts per second (kcps). It reflects the intensity of scattered light and serves as an indirect indicator of particle concentration and sample quality. Data represent mean ± SD from three independent replicates. **(d)** The presence of EVs in pooled fractions 6–12 was confirmed by transmission electron microscopy (TEM; white arrows).

### Proteomic Profiling of Extracellular Vesicles from Boar Reproductive Fluids

Although EVs are small membrane particles, they participate in transporting a large number of proteins. EVs from the caput epididymis contained 6416 proteins, of which only 12 % were unique to this type of EVs. EVs from the corpus epididymis contained the highest number of proteins, namely 6583, but 13 % were unique to this type of EVs. MS identified 5225 proteins from EVs from the cauda epididymis, and only 9 % were unique. The fewest proteins were detected in EVs from SP, specifically 3103 proteins, and 15 % of the proteins were specific only to EVs from SP. On the other hand, 2139 proteins were shared among all four groups of EVs (Figure 5a). The detected proteins in EVs carry a variety of different important functions and are involved in diverse Biological Processes (BPs). Among the top 10 BPs were mainly related to localization, such as cellular, macromolecular, and protein localization, as well as the establishment of this localization. These functions were most specific to epididymosomes. In EVs from SP, BPs included metabolic and catabolic processes in addition to localization processes, and these processes were mainly carbohydrate and protein-related. Transport-related processes, including vesicle-mediated transport, were detected in all four types of EVs (Figure 5b). In the case of KEGG pathways, all four types of EVs contain proteins playing a role in metabolic processes, endocytosis, or the phagosome. EVs from the caput and corpus epididymis contained proteins playing a role in the regulation of the actin cytoskeleton or the proteasome. EVs from the cauda epididymis carried proteins involved in carbon metabolism, oxidative phosphorylation, and glycolysis/gluconeogenesis, which were also detected in EVs from SP. EVs form SP additionally contain proteins that play a role in amino and nucleotide sugar metabolism and biosynthesis or are involved in the biosynthesis of mucin-type O-glycans (Figure 5c). When comparing GO terms between epididymal and seminal plasma samples, the proteins involved in processes within EVs from epididymal fluids are more similar to each other than to those found in EVs from SP.

**Figure 5.**
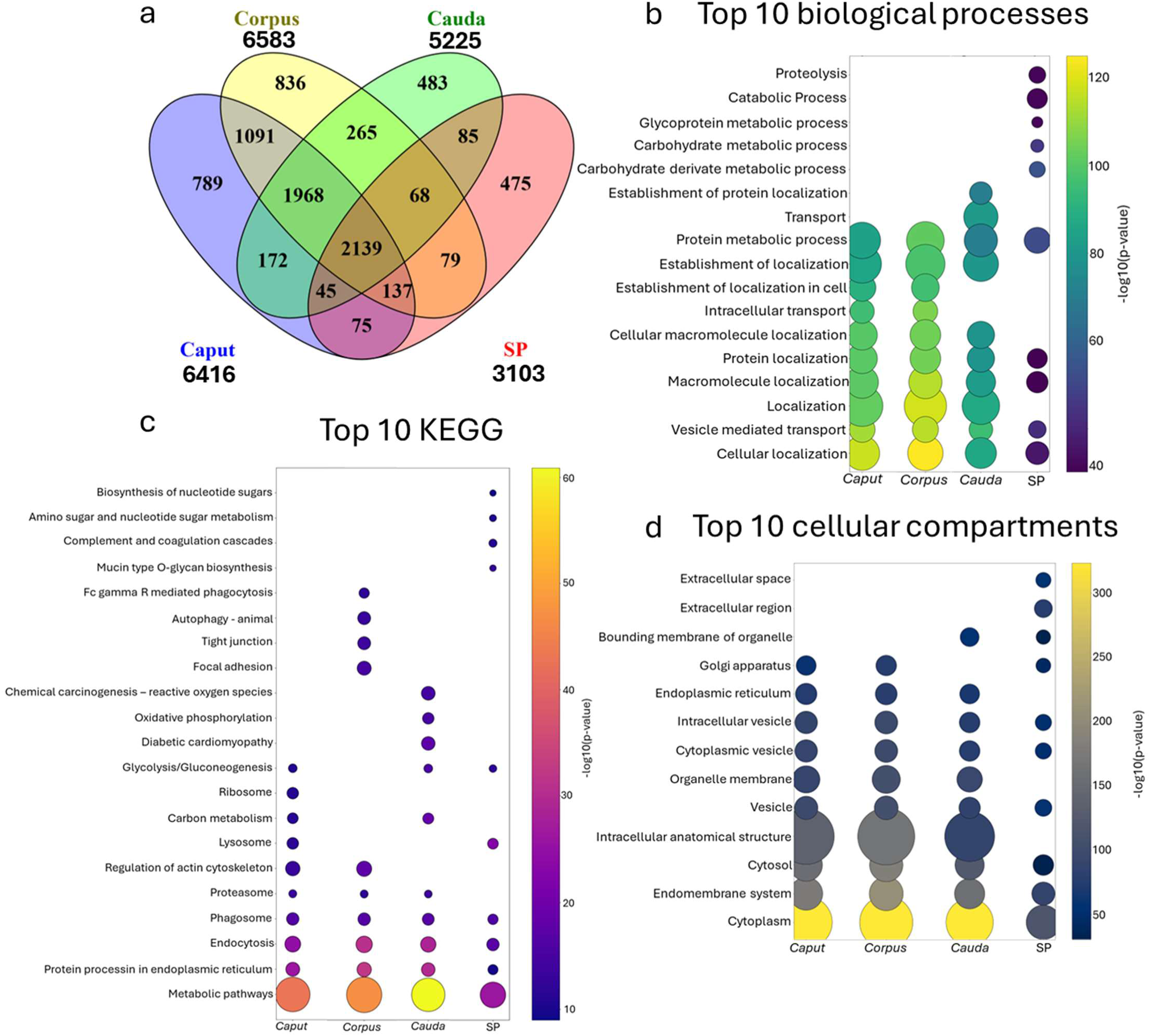
Proteomic analysis of EV cargo. **(a) A** Venn diagram representing the number of identified proteins and their relationships between individual EVs. The numbers inside the closed curves represent the number/percentage representation of a given set. **(b,c,d)** The bubble plots show the top 10 most important biological processes (BPs), Kyoto Encyclopedia of Genes and Genomes (KEGG), and cellular compartments according to GO analysis for proteins of EVs from caput, corpus, cauda epididymis, and seminal plasma (SP). The bubble colour represents the significance (–log₁₀ p-value) of the GO term enrichment (the higher the value, the more significant the result). The bubble size shows the number of proteins from samples that belong to a given GO term (the bigger the number, the larger the bubble).

Besides their functional role, the subcellular localization of proteins identified in EVs is also crucial. Proteins detected in EVs from the male reproductive tract are most localized in the cytoplasm. We also find proteins that are present in the endomembrane system, cytosol, total or cytoplasmic vesicles, or intracellular vesicles. Also, EVs carry membrane proteins, and MS analysis has also found proteins that are specific to the endoplasmic reticulum or Golgi apparatus (Figure 5d).

One of the main markers of EVs is the protein Alix, encoded by the *PDCD6IP* gene. This protein was detected by WB analysis in EVs isolated from all fluids of different parts of the boar reproductive system (Figure 6a). Alix is a protein that is multiply phosphorylated, which affects its molecular weight, resulting in a decrease in its forms from EVs from the caput to SP. While we detected seven forms in the case of caput, the EVs from SP contained only three forms (Figure 6a). MS further confirmed the presence of this protein, and we detected it across a range of molecular sizes in the case of epididymosomes. In EVs from SP, it was only identified by MS in cutting bands of sizes ranging from 100‒50 kDa and from 50‒25 kDa (Supplementary Figure S2). Western blot analysis also revealed fewer ALIX protein bands in SP compared to epididymosomes (Figure 6b). Apart from Alix, MS revealed proteins such as Rab proteins, flotillins, heat shock proteins (Hsp70, Hsp90), or even annexin A2 (ANXA2) (Figure 6c), which play a role in vesicle biosynthesis, transport, or docking. MS also identified proteins that are found in the COPI (Coat Protein Complex I) pathways of vesicles, clathrin-coated vesicles, acrosomal vesicles, and intracellular or extracellular vesicles (Figure 6d).

**Figure 6.**
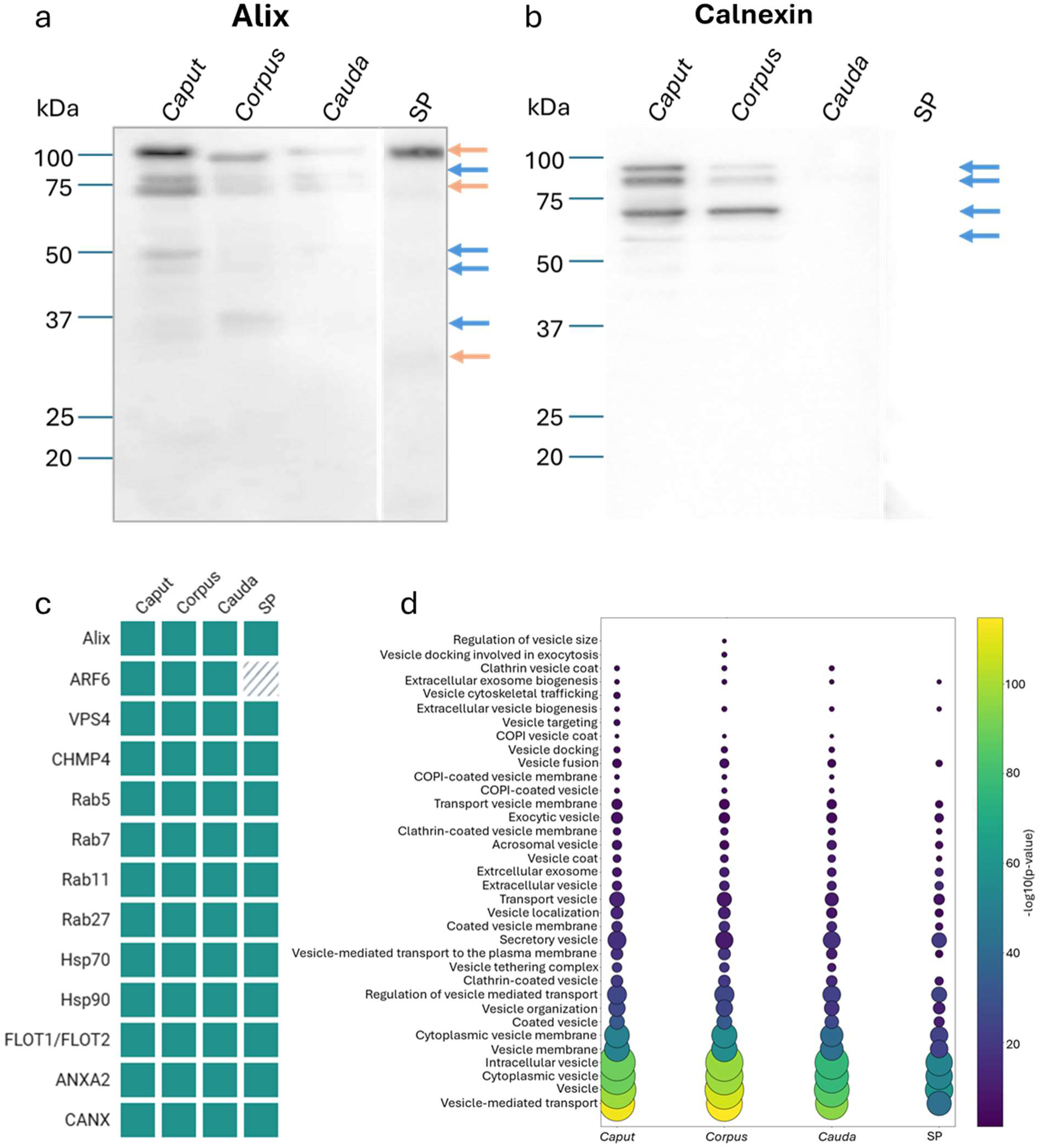
Alix, calnexin, and other proteins play a role in vesicular transport. **(a, b)** Western blot analysis detected Alix and calnexin on EVs from caput, corpus, cauda, and seminal plasma (SP). Blue arrows mark proteins present in epididymosomes, while orange arrows indicate forms specific to both epididymosomes and SP. **(c)** Table of proteins that were detected by mass spectrometry (MS) in EVs isolated from the caput, corpus cauda epididymis, and SP. The green box represents the presence of a given protein in a concrete sample. ARF6-ADP – ribosylation factor 6, VPS4 – Vacuolar Protein Sorting 4, CHMP4 – Charged Multivesicular Body Protein 4, Hsp – Heat shock protein, FLOT – Flotillin, ANXA2 – Annexin A2, CANX – Calnexin. **(d)** Bubble plot representing significantly enriched GO terms connected with vesicles based on proteins identified by MS. Bubble colour indicates statistical significance (–log₁₀ p-value), while bubble size reflects the number of proteins associated with each GO term.

On the other hand, we also observed negative markers of EVs, such as calnexin, a protein of the endoplasmic reticulum. This protein was detected by WB analysis only in epididymosomes from the caput and corpus at different molecular weights (Figure 6b). However, MS identified calnexin also in EVs from SP (Figure 6c).

The following important markers of EVs are proteins from the tetraspanin family – CD9, CD81, CD151, and CD63. We were able to detect the CD9 protein by WB analysis, with one isoform detected at 27 kDa in the case of epididymosomes and two bands detected at 27 and 29 kDa in seminal EVs (Figure 7a). Tetraspanin CD81, as the second marker of EVs, was detected by WB analysis in a similar size to CD9 at 27 and 29 kDa in EVs from SP, but in epididymosomes, the molecular weight was estimated approximately at 18 and 95 kDa (Figure 7b). We were unable to detect CD151 and CD63 tetraspanins using antibodies; however, MS revealed all four tetraspanins in the epididymosomes, and all these proteins, apart from CD9, were identified in EVs from SP (Figure 7c). Tetraspanins are proteins that frequently interact with integrins, which were identified by MS in EVs from the male reproductive fluids (Figure 7c) and play an important role in signalling and adhesion. GO analysis confirmed significant representation of these pathways based on identified proteins. In addition to integrin-mediated signalling and adhesion, pathways including small GTPase-mediated signalling, TOR, HIF-1, AMPK, VEGF, and Rap signalling were also significantly represented by the identified proteins. Also, MS identified proteins involved in hormone-regulated pathways associated with oxytocin or estrogen, as well as those significantly represented in immune, B-cell, and T-cell receptor signalling pathways (Figure 7d).

**Figure 7.**
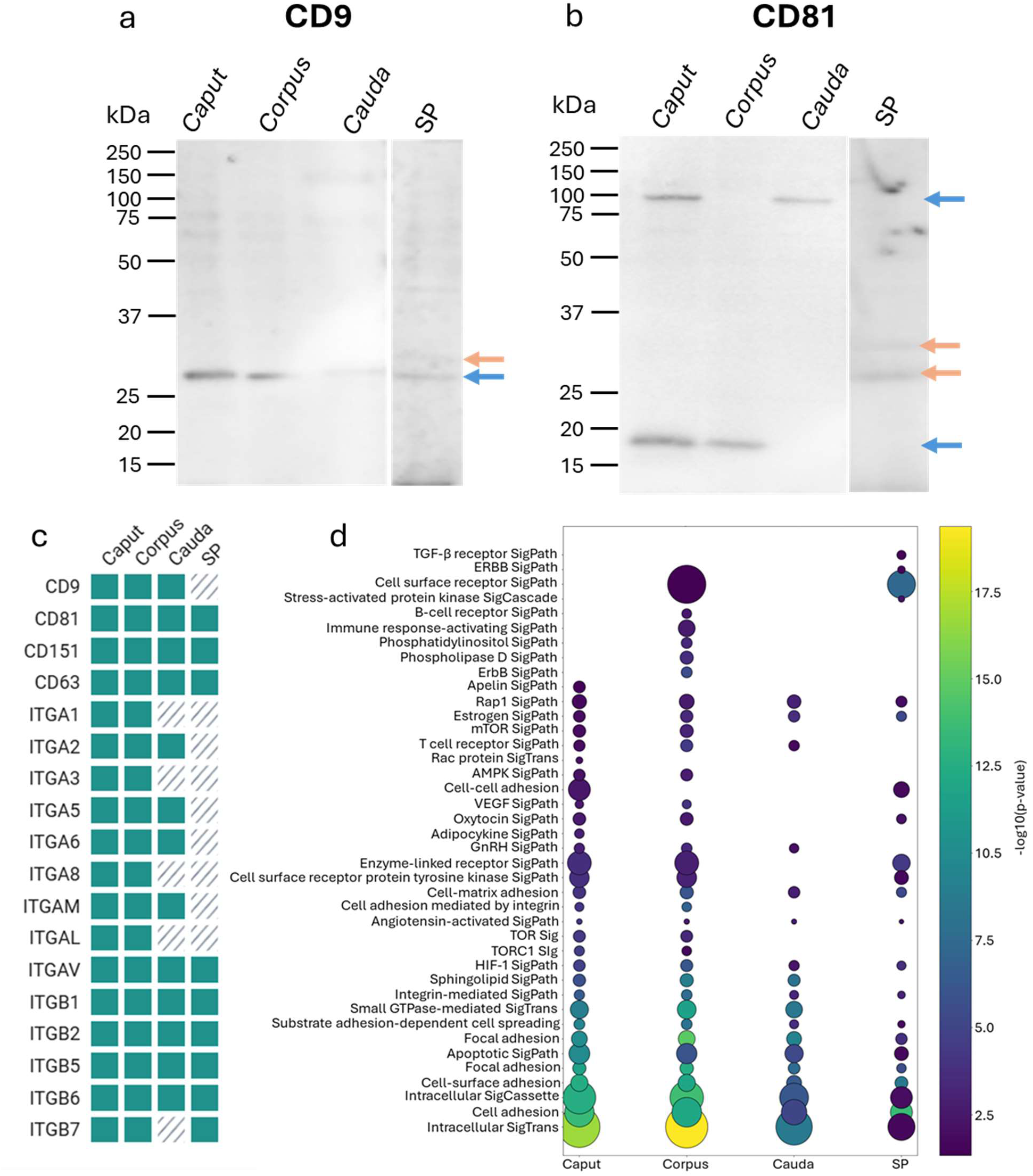
Tetraspanins and integrins detection on EVs. **(a, b)** Western blot analysis confirmed CD9 and CD81 in EVs from caput, corpus, and cauda epididymal fluid and seminal plasma (SP). Blue arrows mark proteins present in epididymosomes and SP, while orange arrows mark forms specific to SP. **(c)** Mass spectrometry (MS) identified tetraspanins and integrins in EVs from the caput, corpus, cauda epididymis, and SP, and their presence is represented by a green box, while a dashed box represents missing protein; ITGA – integrin alpha, ITGB – integrin beta. **(d)** Bubble plot representing significantly enriched GO terms connected with signalling and adhesion based on proteins identified by MS. Bubble colour indicates statistical significance (–log₁₀ p-value), while bubble size reflects the number of proteins associated with each GO term; SigPath – signalling pathway, SigTrans – signalling transduction, SigCassette – signalling cassette, SigCascade – Signalling cascade.

Tetraspanins and integrins play a crucial role in reproduction, working alongside other proteins to ensure the proper course of fertilization. Among such reproductive proteins, MS identified acrosin, ADAM proteins, CD46 protein, clusterin, MFGE8, Rab2A, SPACA, and zonadhesin, which were present in EVs isolated from all four reproductive fluids. Epididymosomes contained A-kinase anchor protein 4 (AKAP4), both spectrin 2 and 4, while spectrin 4 was also present in SP. EVs from the caput fluid contained FCRL5 receptor, while EVs from the corpus were composed of oxytocin receptor (Figure 8a). These proteins indicate that EVs also carry proteins that affect sperm maturation and fertilization processes themselves, which was also confirmed by GO analysis. Among the significant findings were pathways involving proteins that play a role in fertilization itself, sperm-egg recognition, and sperm binding to the zona pellucida, as well as proteins that may be involved in oocyte maturation. The sperm subcellular compartments, in which these proteins are most abundant, are the plasma membrane, flagellum, head, and acrosome (Figure 8b).

**Figure 8.**
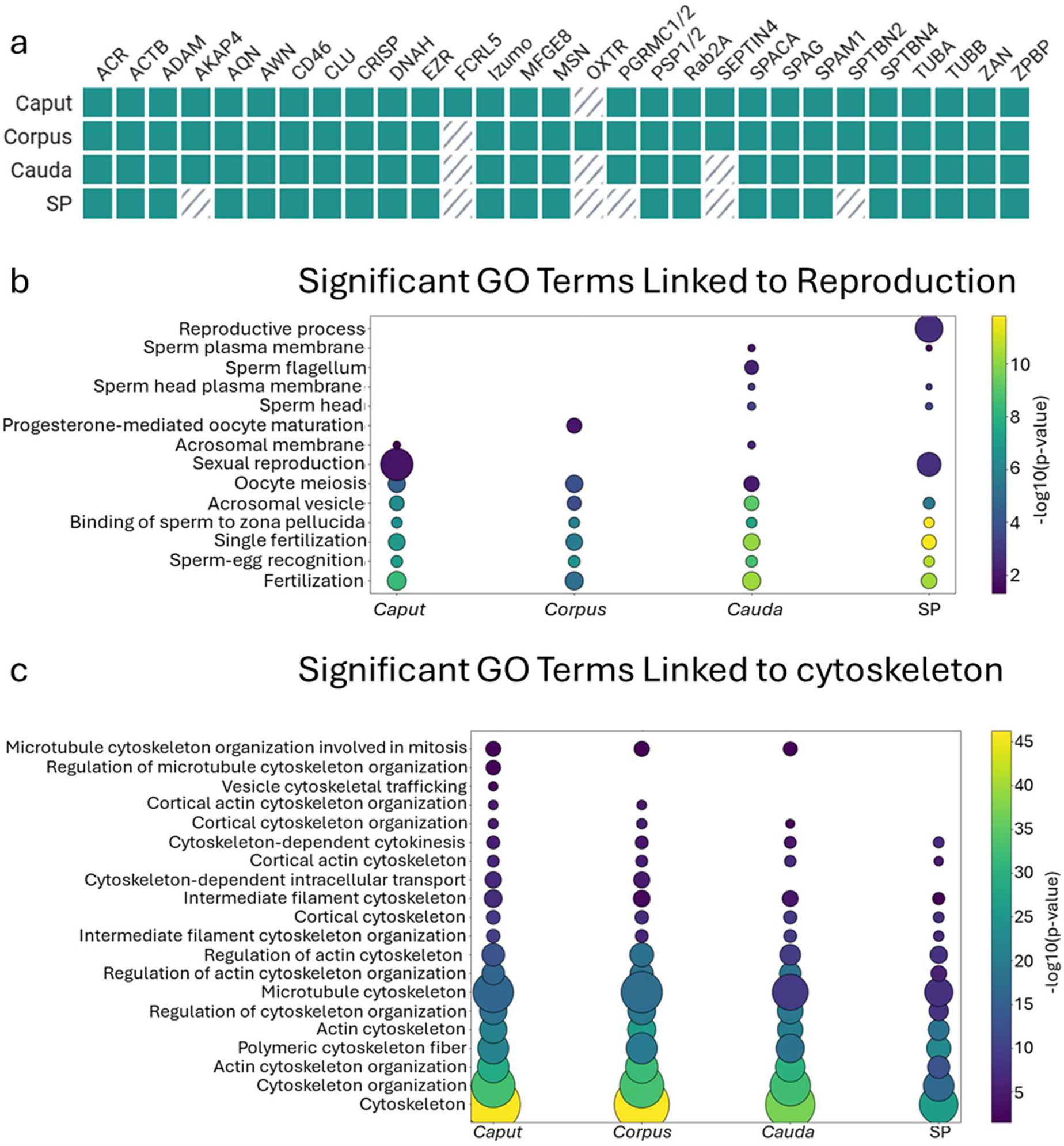
Proteins and pathways in EVs playing a role in reproduction and the cytoskeleton. **(a)** Table of proteins that were detected by mass spectrometry (MS) in EVs isolated from the caput, corpus cauda epididymis, and seminal plasma (SP). The green box represents the presence of a given protein in a concrete sample, while a dashed box represents a missing protein. ACR – acrosin, ACTB – actin, AKAP4 – A-kinase anchoring protein 4, CLU – clusterin, CRISP – cysteine rich secretory protein, DNAH – dynein axonemal heavy chain, EZR – ezrin, FCRL5 – Fc receptor like 5, MFGE8 – milk fat globule EGF and factor V/VIII domain containing, MSN – moesin, OXTR – oxytocin receptor, PGRMC1/2 – membrane-associated progesterone receptor component 1/2, SPACA – sperm acrosome membrane-associated protein, SPAG – sperm associated antigen, SPAM1 – hyaluronidase, SPTBN – spectrin beta chain, TUB – tubulin, ZAN – Zonadhesin, ZPBP – zona pellucida binding protein. **(b,c)** Bubble plots representing significantly enriched GO terms connected with reproduction and cytoskeleton based on proteins identified by MS. Bubble colour indicates statistical significance (–log₁₀ p-value), while bubble size reflects the number of proteins associated with each GO term.

MS analysis also identified proteins that are associated with the cytoskeleton, such as actin, tubulin, as well as ezrin or moesin (Figure 8a). The cytoskeleton, which was confirmed by GO analysis, plays an important role in vesicular transport. GO analysis confirmed several significant pathways based on the proteins identified by MS. These pathways include the cortical cytoskeleton and its organization, as well as cortical actin, intermediate filaments, and regulatory pathways associated with the cytoskeleton (Figure 8c).

### Dynamics of posttranslational modifications of proteins in EVs

Post-translational modifications (PTMs) are an important component of proteins that regulate cellular processes. In EVs, we detected mainly serine phosphorylation of proteins, which had the highest abundance in cauda EVs, but the differences between groups were not significant (Figure 9a,b). Serine phosphorylation, an important PTM in the regulatory pathways of EVs, was observed on several proteins (Figure 5a). Another type of phosphorylation is at tyrosine residues. We were unable to detect this phosphorylation on EV proteins from male reproductive fluids (Supplementary Figure S3); however, MS clearly identified proteins or kinases responsible for phosphorylation of either serine or tyrosine residues (YES, MAPK, JAK1, JAK2, GSK3B), or carbohydrate as well as proteins involved in dephosphorylation in all four groups of EVs (Supplementary Figure S4a,b).

**Figure 9.**
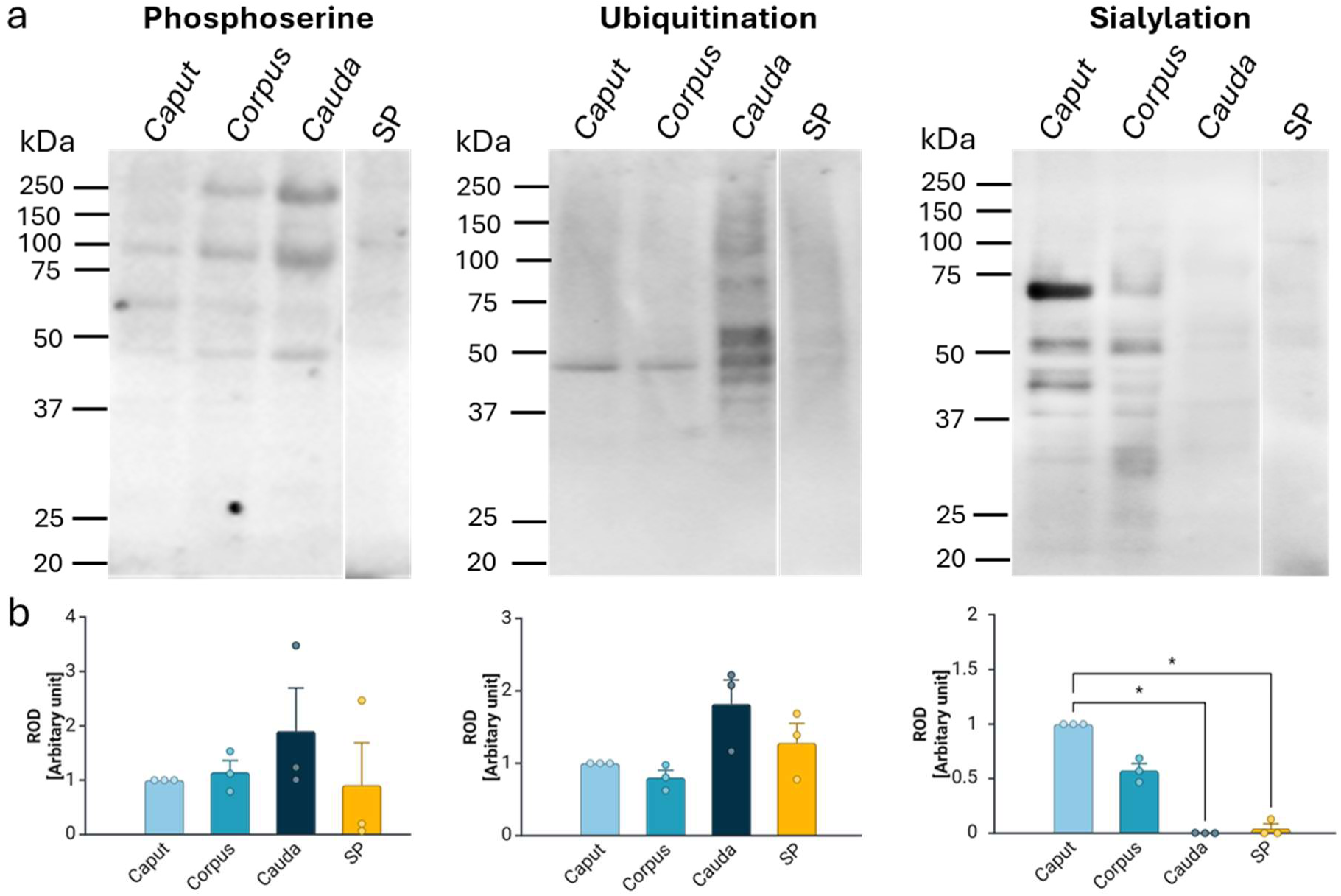
Detection of posttranslational modifications (PTMs) on EVs originated from the epididymis and seminal plasma. (a) Western blot analysis of phosphorylated, ubiquitinated, and sialylated proteins on EVs from caput, corpus, cauda, and seminal plasma (SP). (b) Densitometric analysis of total proteins with PTMs from WB analysis. The Kruskal-Wallis test was used for analysis, and significant results were detected only in the case of protein sialylation from caudal and seminal plasma EVs (p<.05), where a decrease was observed compared to proteins from EVs from the caput epididymis (ROD-relative optical density). Data represent mean ± SEM from three independent replicates.

In addition to phosphorylated proteins, we also detected ubiquitinated proteins, as ubiquitination is one of the pathways involved in protein degradation. Most ubiquitinated proteins were present in EVs from the cauda epididymis and in the SP. However, even in this case, no significant results were observed (Figure 9a,b). Notably, the fact that the highest number of ubiquitinated proteins was found in the cauda and SP EVs also confirms that proteins associated with either ubiquitination or the proteasome system, i.e., protein degradation, were detected in the caput and corpus epididymis, whereas EVs from the SP contained the fewest of these proteins and the associated processes. Among the proteins found in EVs from all four reproductive fluids, CUL3, PSMA2, and UBA1 were identified in MS analysis (Supplementary Figure S4a,b).

The last type of PTMs investigated in this study was protein sialylation, a specific type of primarily terminal glycosylation. We were able to detect sialylated proteins mainly in the caput and corpus. The degree of sialylation in EVs gradually decreased from the caput to the cauda epididymis, while EVs from the cauda epididymis and from the SP contained the least quantity of sialylated proteins. The Kruskal–Wallis test revealed a significant difference among groups, H(3) = 9.84, p = .02. Dunn’s post hoc multiple comparisons indicated a significantly lower value in the cauda compared to the caput (adjusted p = .02) and EVs from SP compared to the caput (adjusted p = .03). No significant differences were observed between the caput and corpus (adjusted p = .89).

Between sialylated proteins, DEFB129 and proteins that participate in sialylation, such as neuraminidase 1 (NEU1), which is responsible for sialic acid (Sia) cleavage, were identified. It was detected in all four groups of EVs of the boar reproductive fluids. On the other hand, transferases, responsible for carbohydrate moieties trafficking, were also found in EVs, but their specificity for binding varied between EVs. Specifically, sialyltransferase profiling revealed that EVs from the caput lacked ST6GALNAC1, those from the corpus did not contain ST6GALNAC2, and caudal EVs did not include ST3GAL1 or ST3GAL6, whereas EVs from the SP were negative for ST8SIA6, among others. Except for proteins responsible for Sia modification, receptors such as SIGLEC were also found among the identified proteins, with SIGLEC1 and SIGLEC10 being specific for EVs isolated from corpus and caput epididymis, respectively. Also, the transporter for Sia-siallin (SLC17A5) was found in epididymosomes. The proteins involved in protein glycosylation were also confirmed by GO analysis, where we determined groups of proteins carried by EVs playing a role in both N- and O-glycosylation, macromolecular glycosylation, or metabolic processes associated with glycosylation (Supplementary Figure S4a,b).

### Functional Crosstalk between EVs and Spermatozoa

In addition to the characterization of EVs themselves, we were also interested in the interaction of EVs with spermatozoa, as well as the transfer of cargos from EVs to spermatozoa. Samples were analyzed by electron and confocal microscopy combined with Imaris visualization, allowing detailed observation of both the localization of EVs on sperm and the transport of their contents.

The interaction of co-incubated EVs with sperm was initially observed by cryo-electron microscopy. EVs originating from SP were clearly bound to spermatozoa from the cauda epididymis, with spherical to oval double-membrane structures (EVs) clearly visible on the sperm head, indicating their preserved morphology and integrity. In the same way, the sperm PM shows no signs of damage or interruption, which excludes the possibility that the observed condition is related to an ongoing acrosomal exocytosis. Furthermore, as shown in Figure 10a, EVs appear to fuse with sperm membrane, providing direct evidence for the transfer of EV cargo to spermatozoa after the membrane interaction (Figure 10a).

**Figure 10.**
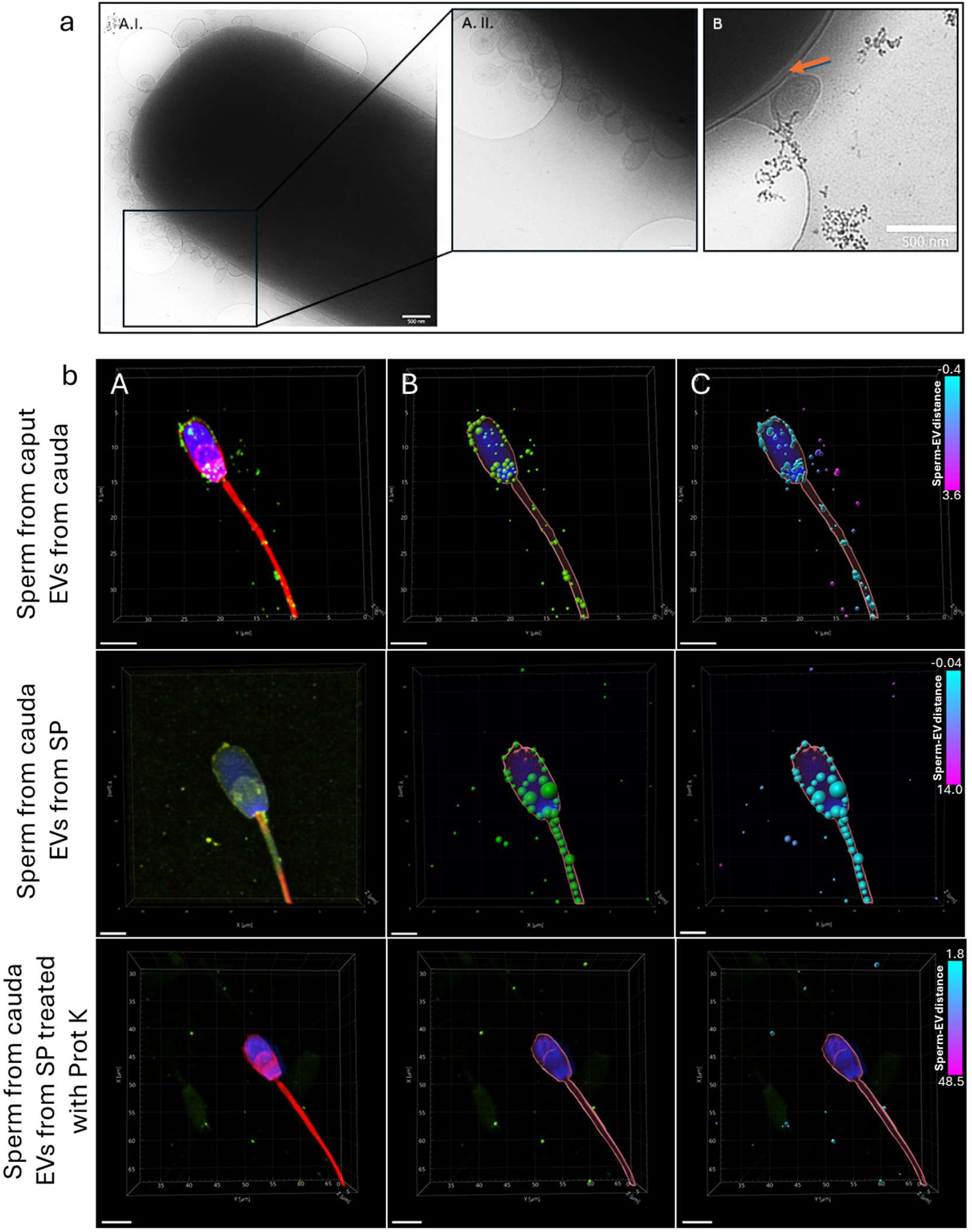
Detection of interaction between the EVs and sperm. **(a)** Detected interaction between the SP-originated EVs and sperm from the cauda epididymis by cryo-electron microscopy. A.I., A.II.) Sperm head with numerous EVs bound to the surface; B) Image showing ongoing membrane fusion between an EV and the sperm cell, with the fusion site indicated by an orange arrow. Scale bar = 500 nm. **(b)** Representative images of sperm with bound EVs. Blue = Nucleus (DAPI), Red = sperm plasma membrane, Green = extracellular vesicles membrane. A) 3D reconstruction of native signal, deconvolved confocal images. B) Imaris processed images with surface reconstruction of sperm membrane and spot representation of EVs. Spot size corresponds to the variable signal intensity of EVs. Nucleus signal is left native. C) Imaris processed images with surface reconstruction of sperm membrane and EVs surface rendering. Colour coding is related to the distance of EVs from the surface of sperm. Nucleus signal is left native. Scale bar = 3 µm.

Sperm-EV interaction was further confirmed by double lipophilic staining, which enabled visualization of EVs’ interaction with the sperm surface and monitoring of binding dynamics using Imaris visualization. Imaris-based 3D image reconstruction and surface modelling (rendering) further illustrated the interaction between EVs and sperm, providing detailed information on EV localization (Figure 10b-B). In the case of cauda epididymal EVs incubated with the sperm from the caput, the strongest signals were detected in the acrosomal and postacrosomal regions and partially along the flagellum. In contrast, SP-originated EVs interacting with the caudal sperm showed signal localization in the flagellum, postacrosomal region, and equatorial segment, with only limited binding in the acrosome. Thanks to color-coded proximity analysis performed in Imaris, EVs showing high signal intensity were confirmed to be in close spatial contact with the sperm membrane, with the possibility of membrane fusion events (Figure 10b-C). As a control, EVs were treated with Prot K, which cleaves surface proteins. Processed EVs were unable to bind to sperm, confirming the importance of surface proteins in the interaction.

Previous experiments confirmed the binding of EVs to sperm and suggested possible membrane fusion. Therefore, we performed experiments for final approval of the EVs fusion with spermatozoa, and cargo release occurs after binding. The following variants of the experimental work were conducted with non-permeable (amine-reactive) and permeable (thiol-reactive) biotin. Non-permeable biotin contains a disulfide spacer arm that prevents this biotin from penetrating the membrane. Therefore, it is ideal for labeling surface proteins. Dot blot analysis showed that biotinylation of native EVs was successful, while EVs treated with Prot K were without signal (Figure 11a). WB analysis of lysed EVs showed biotinylated proteins and their particular molecular weights. In the case of EVs treated with Prot K, no signal was detected (Figure 11b). Co-incubation time-lapse experiments revealed that after incubation of non-permeable biotinylated (NB) EVs with spermatozoa, a positive signal was detected at 30 min, predominantly in the sperm flagellum region (Figure 11c). Any potential effect of EV incubation on spermatozoa was monitored by assessing motility using CASA (Supplementary Figure S5). WB analysis of sperm incubated with NB-treated EVs had no signal in the cytoplasmic protein fraction (Cyt), but this signal was present in the remaining fraction (Rest) of the sperm cells (Figure 11d). However, WB analysis from the time-lapse experiment showed that a biotin signal was detected in sperm at 0, 30, and 60 min, and the most protein bands were present at 60 min (Figure 11d).

**Figure 11.**
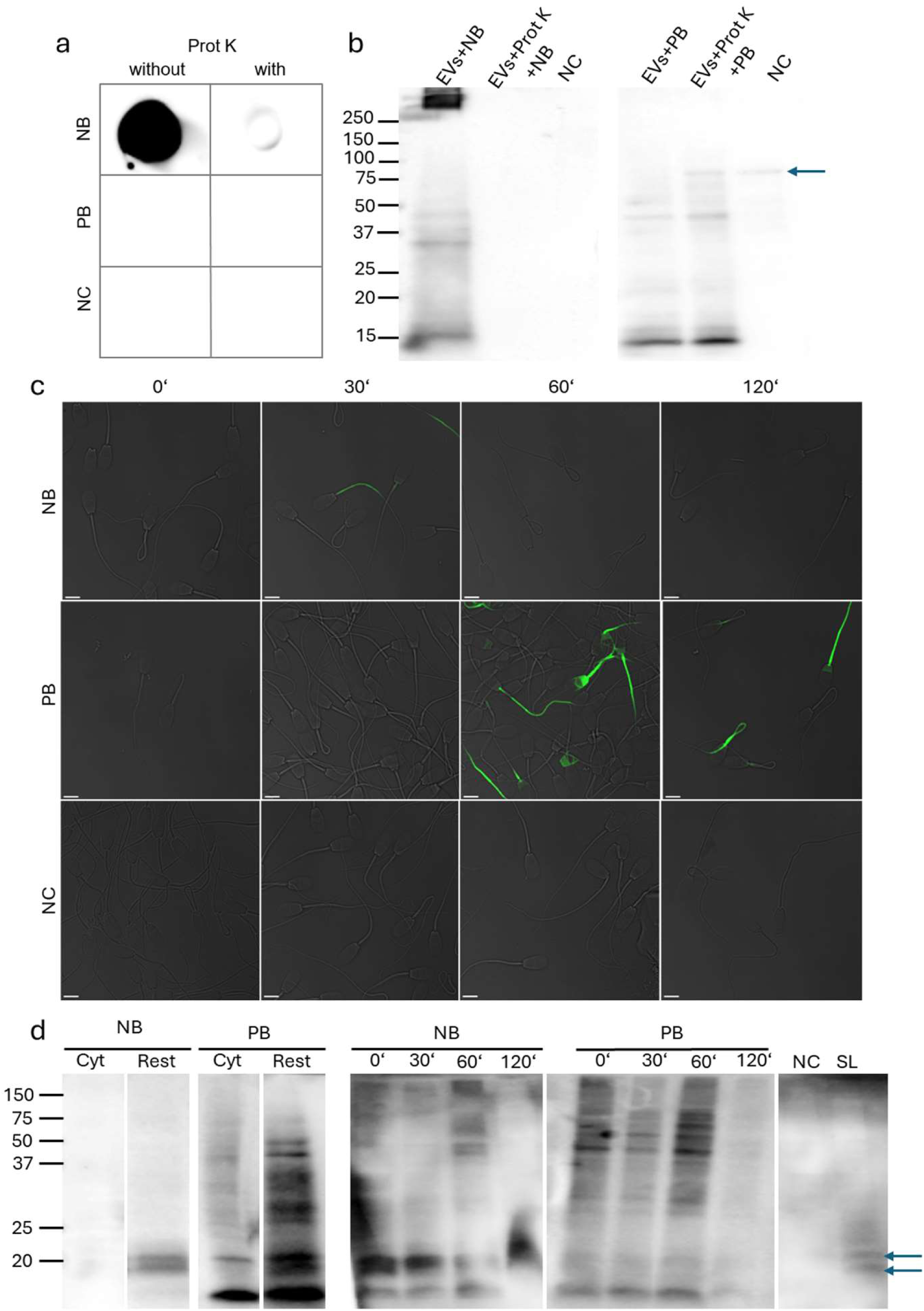
Co-incubating experiments with biotinylation strategy. **(a,b)** Dot blot analysis on native EVs and Western blot (WB) analysis on lysed EVs demonstrated the successful biotinylation of EVs with non-permeant (NB) and permeable (PB) biotin. In contrast, EVs treated with Prot K showed only a weak signal with non-permeant biotin and no signal with permeable biotin. The negative control (NC) represents only EVs without biotin. The blue arrow indicates a nonspecific signal detected also on non-biotinylated EVs. **(c)** Fluorescence microscopy of biotinylated EVs showed that NB (green signal) was detectable at 30 min of the experiment in the tail, while PB (green signal) had the strongest signal at 90 min and a weaker signal at 120 min. NC was without a signal. **(d)** WB analysis of sperm proteins after incubation with EVs showed that NB is not present in proteins from the cytoplasm (Cyt) but is detected in proteins from the remaining part of sperm lysate without the cytoplasmic fraction (Rest), while PB had a detected signal in both Cyt and Rest. Spermatozoa in NB samples/groups showed the strongest signal on WB and the most detected bands at 60 min, while 20 kDa bands were also detected in spermatozoa without EVś co-incubation (SP). Spermatozoa incubated with PB-treated EVs showed a signal at 0, 30, and 60 min. In 120 min, a minimal signal in both NP and PB samples/groups was detected. The blue arrow indicates a non-specific signal after avidin-HRP detection in sperm lysates (SL). The negative control (NC) represents spermatozoa co-incubated with non-biotinylated EVs.

Permeable biotin (PB), containing a stable maleimide group and a non-cleavable cyclohexyl spacer, and due to its membrane permeability, is suitable for permanent labeling of cysteine-containing proteins. However, it does not only label proteins inside EVs but also, to a lesser extent, on the surface, which was confirmed by dot blot analysis, where a weak signal was detected on native EVs (Figure 11a). Lysed EVs also showed a certain amount of signal in EVs treated with Prot K, confirming the presence of biotinylated proteins also inside EVs (Figure 11b). Biotinylated proteins were fluorescently detected in spermatozoa after 60 min of EVś co-incubation, with this signal localization in the flagellum, postacrosomal region, and the equatorial segment. After 120 min, the signal was present only in the sperm flagella (Figure 11c). This demonstrated that not only EVs bind to the sperm membrane but also fuse with it. This result was confirmed by WB analysis (Figure 11d), which detected biotinylated cytoplasmic proteins (Cyt) as well as proteins from the rest sperm fraction (Rest). The time-lapse experiment detected the presence of biotinylated proteins already at time 0 and subsequently at times 30 and 60 min, and to a lower abundance at 120 min. Spermatozoa with non-biotinylated EVs (NC) did not show any signal (Figure 11d).

## Discussion

EVs are important membrane particles that mediate information between cells and play an important role in the reproductive tract, where they ensure communication between sperm and the environment, affect maturation of sperm, immunomodulation, as well as sperm survival in the female reproductive tract, and participate in successful fertilization. They influence all these processes by transporting important cargo containing various ions, metabolites, and macromolecules between cells (Tamessar et al., 2021; Vickram et al., 2021; Roca et al., 2022). Although we have general knowledge about EVs and their role in the reproductive tract, there is a lack of comprehensive information on their mechanisms of action. There are also significant knowledge gaps about EVs between species. While epididymosomes are sufficiently described in bulls, there is minimal information about them in boars. The pig is a very relevant biomedical model organism for humans, and the findings from this model could be applicable to humans in the future. EVs represent a promising therapeutic tool that might be an important part of the improvement of ART for livestock and human clinical practice.

We use for the first time a new optimized method for the isolation of EVs by combining isolation methods via centrifugation, concentration, and size exclusion chromatography (SEC), as mentioned in different formulations in, e.g., the works of Li et al. (2017) and Théry et al. (2018). This approach allowed us to efficiently isolate EVs from all parts of the epididymis (caput, corpus, and cauda) and porcine SP with a significant reduction of contaminants such as cell fragments and free proteins. Ultracentrifugation alone is often associated with low yield, aggregation, or structural deformation of EVs, but a combination of differential centrifugation with SEC can improve the isolation of vesicle-enriched fractions with minimal protein contamination (Lobb et al., 2015; Kamińska et al., 2023). In our study, we aimed to adhere as closely as possible to the MISEV2023 (Minimal Information for Studies of Extracellular Vesicles 2023) guidelines (Welsh et al., 2024) for EVs isolation. However, the process was partially modified by freezing the fluids immediately after initial collection and cell removal. This step, although necessary for sample preservation, may have influenced EVs’ yield, structural integrity, and functional properties, as also evidenced in a small amount of EVs by TEM, which revealed occasional membrane disruptions and altered vesicle morphology. Despite this limitation, the combination and careful optimization of isolation methods enabled us to recover high-quality and functionally competent EVs from epididymal fluids (caput, corpus, and cauda) and SP. These biological fluids contained heterogeneous particles of sizes typically attributed to the exosomes and microvesicles size range (Ståhl et al., 2019). However, given the challenges in distinguishing differences between the different types of EVs based on size or morphology, smaller apoptotic bodies may have been present in the isolated fractions. Although there are markers of EVs such as CD9, CD63, CD81, and TSG101, their presence is not guaranteed in all vesicular populations (Xu et al., 2024a). Apoptotic bodies should be larger compared to other EVs, and, as they are derived from decaying cells, they should contain a different cargo composition, including parts of organelles that are not a regular part of EVs, as described by Poon et al. (2014). However, with ever-evolving research comes the suggestion that apoptotic cells are also likely to produce vesicles comparable in size to microvesicles and exosomes (Kakarla et al., 2020), making both validation of isolation methods and correct interpretation of data difficult. In this context, it is important not only to combine multiple markers of EVs but also to perform complementary functional or morphological analyses that can help to reliably characterize EVs originating from viable cells (Xu et al., 2024a). Nevertheless, our findings support the assumption that boar reproductive fluids contain structurally and functionally diverse EVs, varying in size, morphology, and abundance, with potential region-specific roles. DLS analysis revealed a polydisperse size profile with a dominant population around 100–150 nm, but due to DLS bias toward larger particles and lack of morphological detail, TEM was used to confirm the presence of 50–300 nm vesicles with ‘cup-shaped’ morphology. In comparison with other studies focused on EVs originating from boar SP (Barranco et al., 2019; Parra et al., 2024; Xu et al., 2024b; Xu et al., 2024b), the EVs isolated in this study displayed a similarly heterogeneous size distribution and morphological characteristics. However, EVs from the epididymis were isolated for the first time in this work and therefore could not be assessed in reference to other studies within a single species. Compared to bovine epididymal EVs from the study of Girouard et al. (2011) or mouse epididymosomes described by Barrachina et al. (2022), the epididymal EVs isolated from boar fluids are very similar in size and morphology, but they are less aggregated.

While morphological analysis confirmed that the vesicles have characteristics typical of EVs, proteomic profiling was necessary to reveal their molecular complexity. EVs of the boar reproductive tract are rich in proteins. By comparing epididymosomes with seminal EVs, we detected a lower number of proteins in EVs from SP. We hypothesize that the reason for this is the high heterogeneity, which arises from their diverse tissue origin and distinct lipid composition. Seminal plasma is composed primarily of secretions from accessory sex glands, with each secretion containing EVs released by tissue-specific cells (Schellpeffer and Hunter, 1970; Caballero et al., 2008; Roca et al., 2022). In bulls and mice, differences in lipid composition have been reported between epididymosomes from the caput and cauda epididymis (Rejraji et al., 2006; Girouard et al., 2011). Differences in lipid composition have also been reported in seminal EVs from boars with high versus low sperm motility (Ding et al., 2025). Moreover, the lipid and protein composition is variable between exosomes and microvesicles, which can significantly influence the sufficiency of protein isolation (Haraszti et al., 2016). Based on this, we hypothesize that boar epididymosomes and EVs derived from SP also exhibit such lipid composition differences, potentially affecting protein extraction and downstream analyses. Proteomic profiling further revealed distinct functional signatures across EVs populations. Proteins were clustered into three classes, namely biological processes, KEGG, and cellular localization of proteins. Based on the top 10 detected processes, it is evident that EVs are both heterogeneous and region-specific. The caput epididymis is the initial site of sperm maturation, and EVs originating from this region are enriched in proteins associated with cellular and macromolecular localization, intracellular transport, and processes such as proteasome activity, phagosome function, and protein processing. This protein profile suggests active and dynamic molecular remodelling of proteins occurring in this segment of the epididymis. In addition, pathways associated with metabolism and bioenergetics, such as glycolysis and gluconeogenesis, are represented by proteins in the EVs from the caput. The epididymis is a place that must be highly controlled for quality sperm production (Souza et al., 2017). For this reason, we propose that EVs from the corpus epididymis contain proteins involved in metabolic processes, transport, and localization, as well as proteins associated with tight junction and focal adhesion. These, on the one hand, provide an isolated environment, but on the other hand, they participate in cell communication with the surrounding environment. From the corpus, sperm are delivered to the cauda epididymis, where they are stored and await ejaculation (Rodriguez-Martinez et al., 2022). Therefore, EVs from this part contained proteins not only involved in the processes of localization and transport but also in protein processing, oxidative phosphorylation, glycolysis, gluconeogenesis, and carbon metabolism. These metabolisms are important for maintaining the quality and functionality of spermatozoa, as well as for completing their epididymal maturation (Visconti, 2012; Rodriguez-Martinez et al., 2022; Chen et al., 2023; Badrhan et al., 2024; Simonik et al., 2025). The largest differences in protein clusters are detected for EVs from SP. These proteins are also involved in the transport and localization of EVs and are associated with metabolic processes of sugars, carbohydrates, proteins, and glycoproteins, as well as catabolic processes. This suggests that SP provides sperm with essential macromolecules in the female reproductive tract, which help them survive, protect, and ensure successful fertilization.

Using GO analysis, we determined the typical localization for proteins carried by EVs in the male reproductive tract. These proteins have vesicular, cytoplasmic, cytosolic, or membrane localization. One of the vesicular proteins detected in our samples was Alix, a well-known marker of EVs, which was present in EVs isolated from all four reproductive fluids in a range of molecular sizes, suggesting the presence of different phosphorylated forms, according to the antibody datasheet. This is a very frequent form of PTM for Alix. Interestingly, this PTM regulates its interaction with various protein partners, influencing both its subcellular localization and involvement in diverse cellular processes (Schmidt et al., 2005; Odorizzi, 2006; Sun et al., 2016). In our study, serine phosphorylation was detected as a PTM in EVs across all fluid types, with no significant differences observed between vesicle populations, but a study by Chen et al. (2017) demonstrated differences in phosphorylation between exosomes and microvesicles. Female reproductive EVs transport tyrosine-phosphorylated proteins that may influence, for example, their interaction with sperm or protein stability in the reproductive tract, potentially affecting the acrosome reaction (Fereshteh et al., 2019; Murdica et al., 2020). However, the study by Barranco et al. (2024) demonstrates the opposite, showing that EVs do not affect the tyrosine phosphorylation of sperm or the integrity of the acrosomal membrane. We were unable to detect tyrosine phosphorylation, which is a specific marker of maturation, especially in the capacitation process of sperm. Moreover, we have identified kinases responsible for this type of phosphorylation, and we have also detected GO pathways that are associated with tyrosine phosphorylation. Therefore, we hypothesize that EVs themselves may not only carry phosphotyrosine proteins, but they also transport proteins that are part of the pathways responsible for phosphorylation. These enzymes are activated at the right time on the target cell. On the other hand, protein phosphorylation of EVs may not only be related to their function but also to the loading mechanism. For example, the ARF6-ERK-MLCK (also known as MYLK) pathway participates in the pinching off of microvesicles from the plasma membrane through the phosphorylation of contractile cytoskeletal proteins (Muralidharan-Chari et al., 2009; Szabó-Taylor et al., 2015). ARF6 and MYLK have also been identified as part of the protein profile of epididymosomes in this study. Therefore, EVs may not only transport phosphorylated proteins themselves, but also kinases that can subsequently phosphorylate proteins in target cells. In addition to MYLK proteins, AKT, JAK, MAPK, YES, and SRC kinases have also been identified. Kinases, as regulatory proteins, work closely with small GTPases such as ARF6 or Rab proteins, which ensure proper vesiculation. In addition to ARF6, we have also identified Rab 5, which participates in the formation and sorting of early endosomes (Huotari and Helenius, 2011), Rab7 determining the fate of EVs (Fei et al., 2021, Samuels et al., 2025), Rab11 supporting exocytosis and membrane recycling (Takahashi et al., 2012), and Rab27, which is key to regulating EV formation (Butler et al., 2024). Other proteins that we have identified in epididymosomes and seminal EVs are also closely associated with vesicular transport. VPS4 is connected with the biogenesis of EVs (Jackson et al., 2017), CHMP4 is a very important protein of ESCRT pathways that participates in EVs formation (Hirsova et al., 2016), and Annexin 2 plays a role in vesicular transport and in the docking of EVs (Valapala and Vishwanatha, 2011).

We have also identified proteins associated with the endoplasmic reticulum and Golgi apparatus. A similar observation was reported by Ma et al. (2025), who identified miRNA target genes in seminal EVs. Although proteins derived from intracellular organelles, such as the Golgi apparatus or endoplasmic reticulum, are often considered contaminants in EV samples, their presence may reflect the inherent heterogeneity of EV populations (Welsh et al., 2024). Some studies indicate that “negative markers” may also be present at low levels in certain subpopulations of EVs, particularly during stress or specific cell stimulation (Shahin et al., 2021). Current isolation methods still face limitations in distinguishing between subtypes of extracellular vesicles, including microvesicles and apoptotic bodies, which can carry a diverse range of intracellular proteins. The minimal presence of apoptotic bodies or cell fragments, which are similar in size and sedimentation properties to EVs and can be co-isolated even using the recommended methods, cannot be excluded (Kakarla et al., 2020). Importantly, in the context of the male reproductive tract, this observation gains biological relevance. Spermatozoa are highly specialized cells that are presumably transcriptionally and translationally inactive and lack most organelles, including the endoplasmic reticulum, lysosomes, ribosomes, and Golgi apparatus. Golgi-derived acrosome and the necessity for post-testicular maturation suggest that spermatozoa must rely on external sources, such as EVs, for the acquisition of functional proteins. Therefore, the presence of proteins associated with such organelles in EVs may not merely indicate contamination, which would be a primary interpretation for fluids originating outside the reproductive tract, but rather reflect their role as a molecular reservoir contributing to sperm maturation and function. Such a negative marker is calnexin, a protein of the endoplasmic reticulum (Welsh et al., 2024). We detected this protein in WB analysis in the caput and corpus EVs, but MS detected calnexin in all four types of EVs. Some studies have noted that, despite the different isolation methods used to obtain EVs, calnexin is detected in a population of larger-sized EVs (approximately 200 nm), suggesting that it may be carried explicitly by these EVs (Saludas et al., 2022). This underscores the importance of using multiple markers and analytical methods in validating the purity of isolated EVs and highlights the challenges that must be faced when working with biological material. In addition, the presence of calnexin has been identified as a component of the testicular ADAM2-ADAM3 complex, and association with ADAM2 is crucial for the stability of ADAM3 in epididymal spermatozoa. This complex likely plays a role in sperm interaction with the zona pellucida, suggesting that calnexin may be present in vesicular fractions in association with functionally important protein complexes (Nishimura et al., 2007). We also detected proteins such as histone 3, a nuclear protein, and Golga2, a protein of the Golgi apparatus. In addition, the MS identified proteins that play a role in COPI vesicles, which may also be considered as negative markers of EVs and participate in vesicular trafficking between the Golgi apparatus and the endoplasmic reticulum (Moelleken et al., 2007; Faini et al., 2012). However, we have to emphasize that we detected only proteins and not the compartments themselves, which could be part of EVs and may not have been contaminated solely. By electron microscopy, it was confirmed that the isolated fractions are pure enough. Moreover, the epididymis is a site of sperm maturation and storage. These proteins are likely present precisely due to the proper maturation of spermatozoa. Especially for proteins associated with the Golgi apparatus, since this organelle underlies the formation of the sperm acrosome.

Tetraspanins are also part of the acrosome (Jankovičová et al., 2020a), and these transmembrane proteins were identified among the proteins in our EV fractions. Tetraspanins are not only important for sperm function but also serve as classical markers of EVs (Jankovičová et al., 2020a; Welsh et al., 2024). Using WB analysis, we confirmed the presence of CD9 and CD81, two of the commonly used tetraspanin markers, in agreement with the MISEV2023 guidelines (Welsh et al., 2024). Moreover, MS analysis identified CD9 in epididymosomes and CD81, CD151, and CD63 in EVs from both epididymosomes and seminal EVs. Tetraspanins play essential roles in EV biogenesis, membrane fusion, and interaction with target cells (Andreu and Yáñez-Mó, 2014, Jankovičová et al., 2020b). In addition to their structural role, tetraspanins are involved in cargo selection during exosome biogenesis and may thus influence the functional profile of EVs (Jankovičová et al., 2020b). In reproductive processes, CD9, CD81, CD151, and CD63 have direct or indirect effects on gamete maturation as well as on their interaction and fusion (Jankovičová et al., 2020a). These membrane proteins fulfil their roles largely due to their ability to form tetraspanin-enriched microdomains, which stabilize protein complexes and mediate interactions between cells, e.g., gametes or between cells and EVs (Yáñez-Mó et al., 2009, Perez-Hernandez et al., 2013; Huang et al., 2018; Jankovičová et al., 2020a). Among their most common interaction partners are integrins, formed by α and β subunits, which we identified in our EVs’ samples by MS. The role of integrins, specifically integrin αV, has also been demonstrated in mouse oviductal exosomes, where it participated in the interaction of oviductosomes with sperm (Al-Dossary et al., 2015). Integrin αV was also detected on boar spermatozoa (Palenikova et al., 2021), possibly suggesting that in pigs, integrin αV, along with tetraspanins, may mediate the interaction of epididymosomes and seminal EVs with spermatozoa. Integrins are known to mediate both cell adhesion and intracellular signalling (Merc et al., 2021). MS identified proteins as part of EVs that are significantly represented in intracellular signalling and transduction, cell and focal adhesion, as well as integrin-mediated signaling. Moreover, we also detected proteins involved in key signaling pathways, such as the TOR (Target of Rapamycin) pathway, which is responsible for protein phosphorylation (Chiang and Abraham, 2005), and pathways mediated by small GTPases, further supporting the involvement of these vesicles in dynamic intercellular communication. We also detected signalling pathways associated with hormones such as oxytocin and estrogen, as well as those connected to the immune response. The immune response must be tightly regulated in both the male and female reproductive tract (Nguyen et al., 2014), and EVs are very likely involved in these processes. Another way EVs can participate in modulating the immune response is through sialylation. Sia, as a component of glycoconjugates, can participate in modulating the immune response via SIGLEC receptors (Zhu et al., 2024). EVs from the corpus transferred receptors SIGLEC1 and SIGLEC10 were identified in EVs from the caput. It is assumed that Sia masks important antigens on sperm for the immune system (Tecle and Gagneux, 2015). It has been shown in humans that higher sialylation of SP proteins is associated with increased levels of anti-sperm antibodies and decreased sperm quality (Palenikova et al., 2024). We detected sialoproteins as part of EVs, mainly from the caput and corpus epididymis, where they apparently were significant for sperm maturation and plasma membrane remodelling, whereas EVs from the cauda and SP exhibited reduced sialylation. In this study, we focused on α-2,6-linked Sia; while other types of Sia linkage may also be present in these EVs, their detection requires further analysis. Among the sialylated proteins, we identified DEFB129, which belongs to the beta-defensin family. There are small anti-microbial peptides that are responsible for defence against pathogens such as *E. Coli* (Zeng et al., 2022). DEFB129 is in the same family as DEFB126, which has been confirmed as a sialylated protein responsible for sperm transport through the acidic environment of the female reproductive tract (Tollner et al., 2008, Tollner et al., 2012). In addition, we detected enzymes responsible for Sia modification, specifically neuraminidase 1, which cleaves Sia and was present in all EVs from the boar reproductive tract. Transferases involved in Sia binding were detected more specifically. For example, ST3GAL1 and ST3GAL6 were not present in EVs from the cauda; STLGALNAC1 was not present in EVs from the caput; ST6GALNAC2 was not present in EVs from the corpus and cauda; while ST6GALAC4 was identified in all four types of EVs. On the other hand, EVs from the SP did not contain sialyltransferase ST8SIA6, which was only identified in EVs from the epididymis. Therefore, it is possible that EVs themselves do not specifically transfer sialoproteins to sperm but rather transfer the enzymatic machinery that ensures the proper processing of glycoconjugates. Proteins responsible for overall N- and O-glycosylation had the highest representation in EVs from SP according to GO analysis. Glycoconjugates, especially sialylation, are vital for sperm function and fertilization. Forming the glycocalyx, they support sperm passage through the female tract, mediate immune protection, and enable interactions with oviductal cells. Sialylation specifically regulates sperm surface remodeling and helps delay capacitation, ensuring proper timing for fertilization (Tecle and Gagneux, 2015).

Another PTM that we successfully detected is ubiquitination in proteins isolated from EVs. Its exact role is unknown, but ubiquitination is an important PTM mainly in the process of protein degradation by the proteasome and is crucial for sperm maturation and quality control (Sutovsky, 2003). We have identified proteins that play both a role in ubiquitination and are also part of catabolic pathways. While it remains unclear whether these catabolic pathways are activated directly within EVs or only upon delivery to target cells, it is evident that ubiquitination is an important protein PTM in EVs. It is also interesting that most of these pathways are present in EVs from the caput and corpus epididymis. We hypothesize that this reflects the role of the caput in initiating sperm maturation (Cornwall, 2009, Dacheux and Dacheux, 2014), where regulation via ubiquitination and related catabolic processes is particularly required. Cauda EVs show the highest levels of ubiquitinated proteins. However, this modification may already be initiated in the preceding regions of the epididymis, where proteins responsible for ubiquitination are more abundant. Moreover, it is possible that if ejaculation does not occur in time, these ubiquitinated proteins are the ones that are doomed to degrade from damaged, dead, and poor-quality sperm. However, ubiquitination may not only play a role in degradation but may also influence the cargo of EVs. For example, the E3 ligase TRIM25, which we also identified using MS in epididymosomes, participates in K-63-linked ubiquitination of the FMR1 protein, which influences its loading into inflammation-mediated EVs (King et al., 2023). Therefore, further experiments should clarify the type of ubiquitination and whether it mediates protein degradation or is crucial for cargo sorting.

To further explore the EV cargo sorting, we investigated the interaction between EVs and sperm and monitored the transfer of EV cargo. Using various microscopic techniques, we demonstrated that EVs can actively bind to and deliver their contents into sperm cells through dual lipophilic and biotin-based labeling strategies. Firstly, cryo-electron microscopy captured the interaction between the EVs and sperm, as well as their possible membrane fusion as the mechanism for transferring the vesicular cargo into sperm. The interaction between EVs and sperm was further supported by dual lipophilic labeling, which enabled the visualization of EV binding, specifically, EVs from the cauda binding to sperm from the caput and seminal EVs interacting with cauda sperm mimicking processes during sperm maturation in the epididymis and events at ejaculation. Using Imaris post-processing image analysis, we were the first to obtain a detailed 3D visualization of these interactions, showing that cauda EVs primarily associate with the acrosomal and postacrosomal regions of the caput sperm, with some apparent binding also along the flagellum. This spatial pattern may reflect the functional specialization of the cauda epididymis, which is primarily responsible for storing and maintaining the viability and motility of sperm during this phase. It is therefore plausible that EVs deliver essential nutrients or regulatory proteins to support sperm maintenance and survival (James et al., 2020, Sullivan, 2015). Proteomic analysis revealed specific protein cargo within the epididymosomes. Proteins such as AKAP4, PGRMC1/2, and SPTBN2 were consistently identified across all types of epididymosomes, suggesting their broad involvement in sperm maturation. In contrast, FCRL5 was uniquely present in caput-derived EVs, OXTR was specific to corpus-derived EVs, and SEPTIN4 was found in both caput and corpus EVs. Interestingly, several proteins, including ACR, members of the ADAM family, CD46, CLU, tubulins, ZAN, and Rab2A, were detected across all four EVs’ populations, indicating their potential roles in EV-mediated sperm modulation. These proteins primarily serve as cargo for sperm maturation or play direct roles in mediating EVs-sperm membrane fusion, analogous to the function of Izumo or tetraspanins during gamete interaction (Inoue et al., 2005; Jankovičová et al., 2020a; Al-Dossary et al., 2015). There is an open question remaining for further investigation. However, Zhou et al. (2024), using a mouse model, suggested that the fusion of epididymosome membranes with sperm depends on the integrity of lipid rafts and the presence of dynamin 1, which was also identified by MS in isolated EVs. This highlights the importance of investigating the molecular mechanisms underlying EV-cell interactions with an emphasis on detailed analysis of PM characteristics. One of the candidate proteins identified as a regulator of the interaction between sperm and EVs from SP was ezrin (Xu et al., 2024b), membrane-cytoskeleton linker protein, which we also recognized as a part of EVs proteome using MS. SP contains components including EVs that ensure the sperm motility and their membrane integrity, antioxidant capacities, surviving sperm in female reproductive tract and helping to time capacitation using decapacitation factors such as spermadhesins (Samanta et al., 2018, Du et al., 2016), which was identified as part of EVs in this study.

In addition to demonstrating the interaction between EVs and sperm, we were also able to capture the transfer of cargo between EVs and sperm using both the permeable and impermeable biotin. Zhou et al. (2019) have already demonstrated in a mouse model that the initial interaction between EVs and sperm occurs predominantly in the postacrosomal region. Our results are consistent with these findings, as dual staining and biotinylation confirmed a similar localization of EVs binding, followed by cargo release, which was most evident at 60 minutes post-incubation. This was particularly evident in the experimental group with intra-EVs cargo labeled by permeabilization strategy. Unlike the results of Zhou et al. (2019), a gradual time-dependent increase in cargo signal is observed. We hypothesize that this discrepancy may be attributed to the use of a different animal model or a different experimental procedure. Our data clearly demonstrate not only the physical interaction between EVs and sperm but also the successful transfer of EV cargo, both external and internal, into sperm cells.

## Conclusion

This study provides the first comprehensive characterization of epididymosomes in pigs, analyzing EVs from the caput, corpus, cauda epididymis, and seminal plasma. Precise proteomic profiling revealed classical vesicular markers (Alix, tetraspanins) alongside proteins involved in signalling, immune modulation, and reproduction. We also identified phosphorylated, ubiquitinated, and sialylated proteins, with subtype-specific patterns, and enzymes likely involved in PTMs. Importantly, we demonstrated not only the interaction of EVs with sperm but also the transfer of EV-derived cargo into sperm cells. Despite certain limitations, our findings significantly advance the understanding of EV-mediated communication in the male reproductive tract and the potential role of EVs in sperm maturation. Moreover, given the physiological similarities between pigs and humans, insights gained from this study may also advance our understanding of reproductive processes in humans and support the development of clinical applications.

## Author Contributions

**Veronika Kraus** design of the study, methodology, experimental work, image and data analysis, result interpretation, original draft and writing of the manuscript; **Barbora Doleckova** design of the study, methodology, experimental work, result interpretation, original draft and writing of the manuscript; **Ondrej Sanovec** imaging, Imaris analysis; **Michaela Frolikova** imaging; **Daniela Spevakova** Western blot analysis; **Aneta Pilsova**, **Zuzana Pilsova** experimental work; **Katerina Komrskova** manuscript editing, funding acquisition; **Ondrej Simonik** design of the study, data analysis, supervision, original draft and writing of the manuscript; **Pavla Postlerova** design of the study, methodology, data analysis, result interpretation, original draft and writing of the manuscript, supervision, funding acquisition. All authors contributed to the manuscript editing and approved the final version.

## Funding

This study was supported by the Czech Science Foundation GA22-31156S, the Ministry of Education, Youth and Sports of the Czech Republic under the INTER-EXCELLENCE II program, subprogram INTER-ACTION LUAUS25072, the Internal Grant Agency of the Czech University of Life Sciences in Prague SV24-21-21230 and SV25-2-21230, project Andronet (CA20119) supported by COST, Research programme Strategy AV21 Future of Assisted Reproduction (ART), and by the institutional support from the Institute of Biotechnology RVO: 86652036.

## Acknowledgements

We acknowledge Jan Rasl from the Structural Mass Spectrometry Core Facility, CF of Biophysics of CIISB, Instruct-CZ Centre, supported by MEYS CR (LM2023042) and European Regional Development Fund-Project „UP CIISB“ (No. CZ.02.1.01/0.0/0.0/18_046/0015974). The authors also acknowledge Jiří Mikšátko and David Liebl from the Imaging Methods Core Facility at BIOCEV, supported by the MEYS CR (LM2023050 Czech-BioImaging), for their support & assistance in this work, and the Imaging Methods Core Facility of the BIOCEV, the Light Microscopy Core Facility, IMG, Prague, CR, supported by MEYS (LM2023050, CZ.02.1.01/0.0/0.0/18_046/0016045, and CZ.02.01.01/00/23_015/0008205). The authors thank Biofarm Sasov for providing boar reproductive tissues, and the Skrsin insemination station (Lipra Pork, a.s., Rovensko pod Troskami, Czech Republic) for supplying boar semen.

## Conflicts of Interest

The authors declare no conflict of interest.

## Supplementary materials

### Proteomic Profiling of Extracellular Vesicles from Boar Reproductive Fluids

**Supplementary Figure S1.**
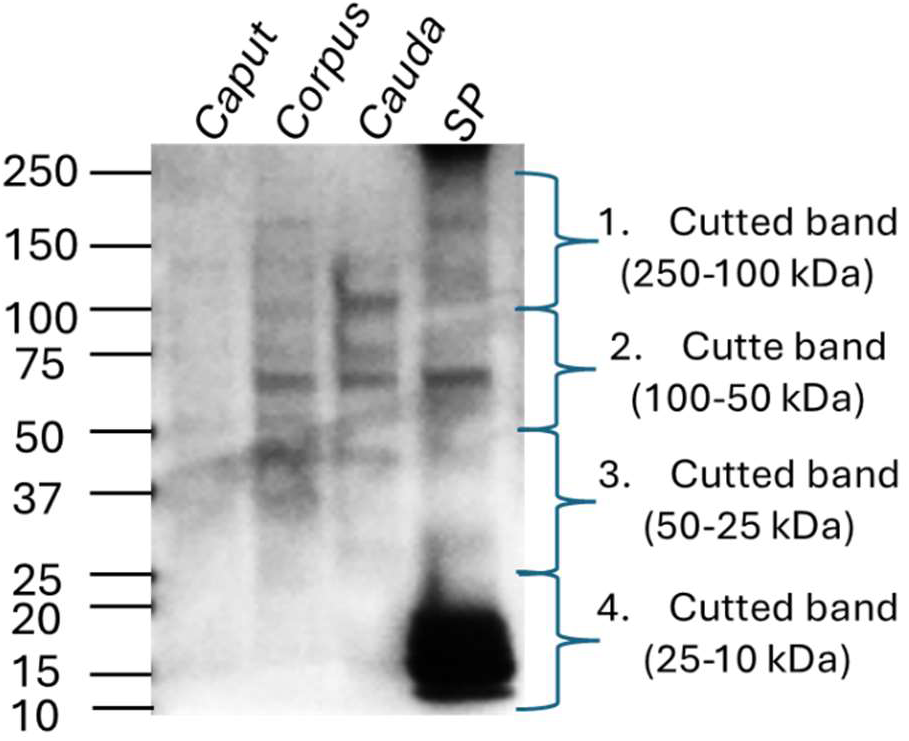
Protein staining in gel by Coomassie brilliant blue solution. Four protein bands were cut from the gel, corresponding to molecular weights of 250‒100 kDa, 100‒50 kDa, 50‒25 kDa, and 25‒10 kDa. These protein bands were analyzed by mass spectrometry.

**Supplementary Figure S2.**
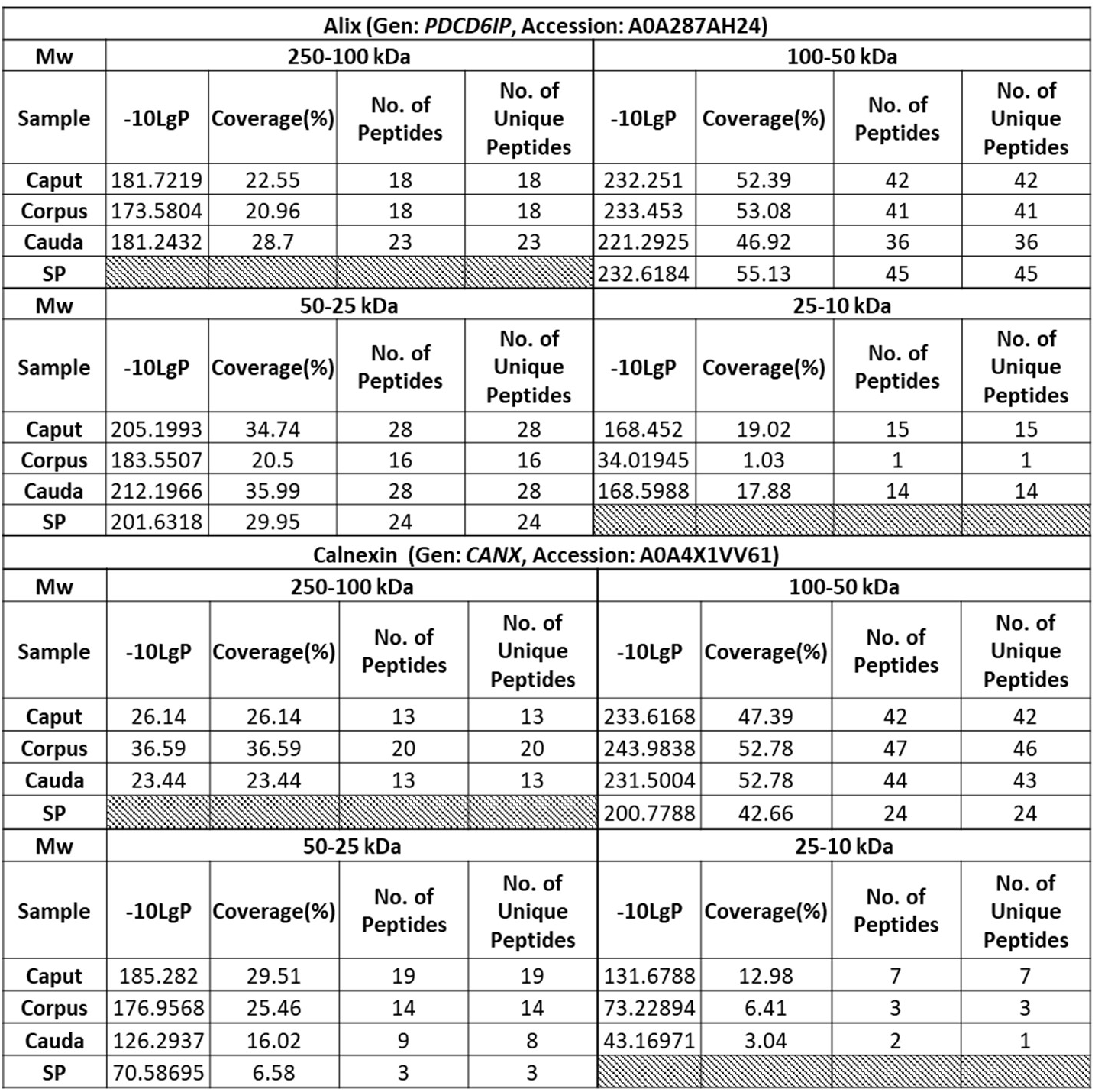
Identification of proteins Alix and calnexin in different molecular weights (Mw) by mass spectrometry (MS). EVs from caput, corpus, and cauda epididymis contained Alix and calnexin in all four groups of Mw that were subjected to MS, while EVs from seminal plasma (SP) contained these proteins only in bands cut from the gels of molecular weights 100‒50 kDa and 50‒25 kDa. The -10LgP (minus ten times the logarithm (base10) of the P-value) represents the probability that the identified protein is a random match, Coverage (%) represents the percentage of protein covered by identified peptides, No. of Peptides is the number of identified peptides by MS, and No. of Unique Peptides is the number of peptides unique to specific proteins.

### Dynamics of Posttranslational Modifications of Proteins in EVs

**Supplementary Figure S3.**
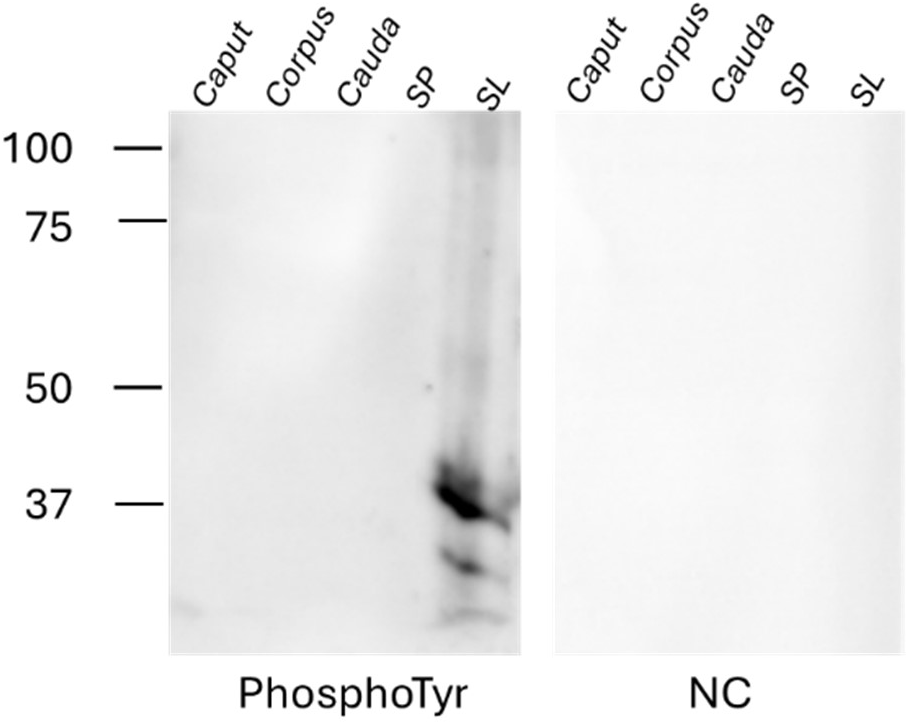
Detection of proteins phosphorylated on tyrosine. The phosphorylated proteins on tyrosine were detected only in the sperm lysate (SL), and EVs from caput, corpus and cauda epididymis, and SP are without any specific signal. Negative control (NC) was incubated without primary antibody.

**Supplementary Figure S4.**
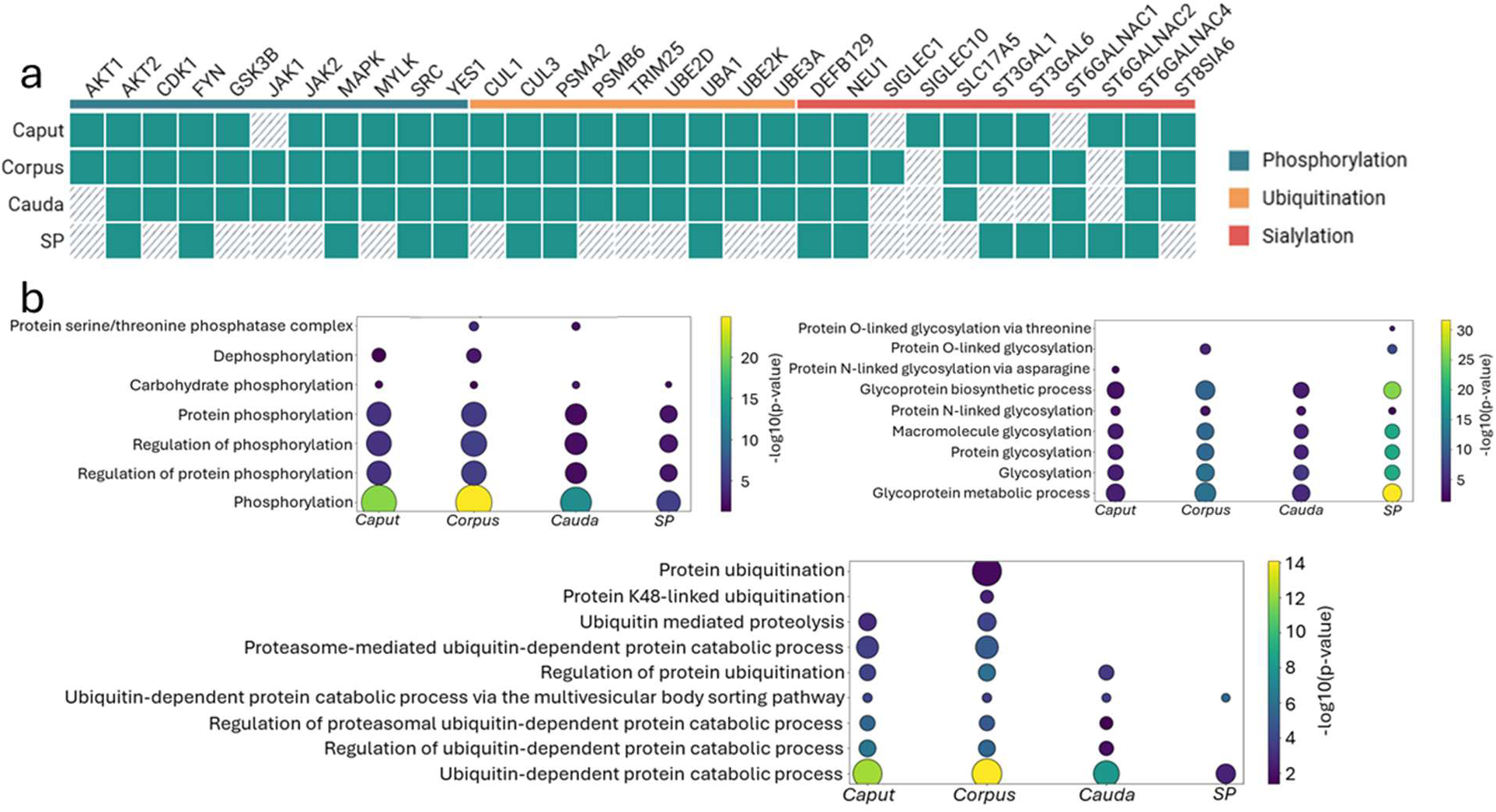
GO analysis of proteins connected with PTMs. (a) Table of proteins detected in EVs isolated from the caput, corpus cauda epididymis, and seminal plasma (SP) by mass spectrometry (MS). The green box represents the presence of a given protein in a specific sample, while the dashed box represents a missing protein.

Tyrosine – protein kinase YES or SRC; Mitogen-activated protein kinase – MAPK; Tyrosine-protein kinase JAK2, JAK1, FYN; Glycogen synthase kinase-3 beta – GSK3B; Cell division cycle 2 – CDK1; non-specific serine/threonine protein kinase – AKT2, RAC-alpha serine/threonine protein kinase – AKT1; Cullin – CUL; Proteasome 20S subunit alpha 2 – PSMA2; Proteasome 20S subunit beta 6 – PSMB6; E1 ubiquitin-activating enzyme – UBA1; Ubiquitin conjugating enzyme E2 D – UBE2D, Ubiquitin conjugating enzyme E2 K – UBE2K; Ubiquitin-protein ligase E3A – U BE3A; Beta-defensin 129 – DEFB129; Neuroaminidase – NEU1; Ig-like domain-containing protein – SIGLEC; Sialin – SLC17A5,CMP-N-acetylneuraminate-beta-galactosamide-alpha-2,3-sialyltransferase 1 – ST3GAL1; Type 2 lactosamine alpha-2,3-sialyltransferase – ST3GAL6; alpha-N-acetylgalactosaminide alpha-2,6-sialyltransferase – ST6GALNAC1; alpha-N-acetylgalactosaminide alpha-2,6-sialyltransferase – ST6GALNAC2; N-acetylgalactosaminide alpha-2,6-sialyltransferase 4 – ST6GALNAC4; Alpha-2,8-sialyltransferase – ST8SIA6. (b) Bubble plot representing significantly enriched GO terms connected with phosphorylation or dephosphorylation, ubiquitination, or glycosylation of proteins.

### Functional Crosstalk between EVs and Spermatozoa

**Supplementary Figure S5.**
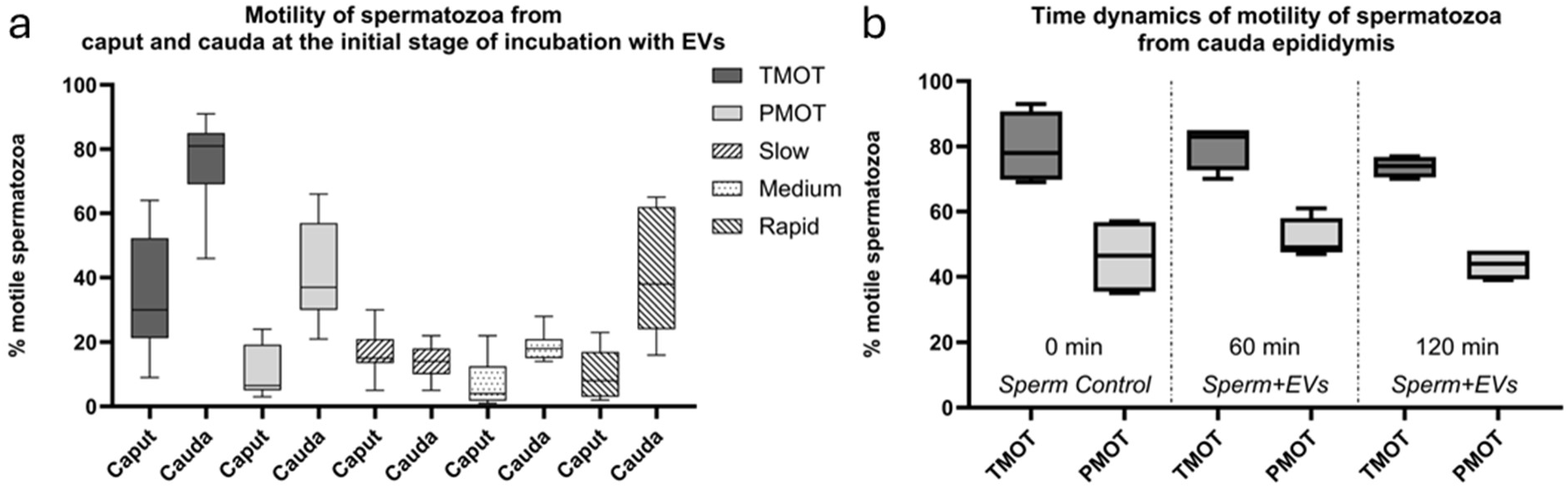
Motility of sperm. (a) At the initial stage of the experiment, the motility of spermatozoa isolated from the caput and cauda epididymis was evaluated, confirming their initial quality and functional competence. (b) During the biotinylated experiment, the motility of caudal spermatozoa incubated with EVs was monitored over time, showing no significant (p>.05) changes in motility or overall sperm quality throughout the incubation period.

## Notes

### Competing Interest Statement

The authors have declared no competing interest.

